# Promoting axon regeneration by enhancing the non-coding function of the injury-responsive coding gene *Gpr151*

**DOI:** 10.1101/2021.02.19.431965

**Authors:** Bohm Lee, Jinyoung Lee, Yewon Jeon, Eunsoo Jang, Yeonsoo Oh, Hyemin Kim, Minjae Kwon, Jung Eun Shin, Yongcheol Cho

**Affiliations:** Department of Life Sciences, Korea University, Seoul 02841, Republic of Korea; Department of Molecular Neuroscience Dong-A University College of Medicine, Busan 49201, Republic of Korea; Department of Translational Biomedical Sciences, Graduate School of Dong-A University, Busan 49201, Republic of Korea

**Keywords:** 5’UTR, axotomy, axon regeneration, RNA-binding protein, translation, *GPR151*, single-exon gene, CSDE1, dorsal root ganglion, *RiboTag*

## Abstract

Gene expression profiling in response to nerve injury has been mainly focused on protein functions of coding genes to understand mechanisms of axon regeneration and to identify targets of potential therapeutics for nerve repair. However, the protein functions of several highly injury-induced genes including *Gpr151* for regulating the regenerative ability remain unclear. Here we present an alternative approach focused on non-coding functions of the coding genes, which led to the identification of the non-coding function of *Gpr151* RNA interacting with RNA- binding proteins such as CSDE1. *Gpr151* promotes axon regeneration by the function of its 5’- untranslated region (5’UTR) and expression of an engineered form of the 5’UTR improves regenerative capacity in vitro and in vivo in both sciatic nerve and optic nerve injury models. Our data suggest that searching injury-induced coding genes potentially functioning by their non- coding regions is required for the RNA-based gene therapy for improving axon regeneration.

## Introduction

Neurons in the peripheral nervous system (PNS) activate the intrinsic regeneration program after axons are injured, a process that is controlled by the expression of injury- responsive genes^1–6^. Comparative analysis of transcriptomic data between naïve and regenerating sensory neurons in dorsal root ganglia (DRG) has revealed a number of differentially expressed genes (DEGs), within which there is a subgroup of regeneration- associated genes that are required for successful regeneration ^7–9^. However, the molecular mechanisms of the particular regeneration-associated genes involved in the regulation of regenerative potential have not been clearly delineated.

Functional analysis of protein-coding DEGs is generally regarded as a reliable strategy for understanding cellular mechanisms. This paradigm requires proof that upregulated protein- coding mRNA is directed to ribosome complexes if it is believed that the function of DEGs results primarily from the resulting proteins. However, with respect to the injury-responsive upregulated genes (IRGs), several reports have shown a poor correlation between their mRNA and protein levels^10–13^, indicating that it is necessary to study the differentially expressed mRNA without the assumption that their function as regeneration-associated genes is dependent on their protein products. Therefore, monitoring the fate of the mRNA itself may be the primary route to correctly understanding how neurons alter their physiology to enter a regenerative state via DEGs^14–17^.

RNA-binding proteins (RBPs) play various roles including the regulation of RNA for its stability, translational efficiency, modification, localization and intermolecular interaction ^18, 19^. RBPs such as the CELF family protein UNC-75 affect axon regeneration in *C. elegans* via regulating alternative splicing and the Zipcode-binding protein ZBP1 can promote nerve regeneration in mice by regulating axonal mRNA transport^20, 21^. Recent studies have shown that mRNA differentially expressed by specific cues can be directed to RBPs and modulate their regulatory functions^22, 23^. These reports raise a hypothesis that the injury-dependent mRNA may affect the regenerative program by regulating RBPs independently from the function of its translated protein.

In this study, we conducted a comparative analysis of transcriptomes under an axon regeneration paradigm based on their ribosome-association efficiency using *RiboTag* mice after axonal injury^24^. We found that a group of injury-responsive genes exhibited no increase in their ribosome-association efficiency even when their mRNA was dramatically upregulated after axonal injury, supporting potential non-coding functions of these mRNA. Of these genes, *Gpr151* encodes an orphan G protein-coupled receptor (GPCR) that is implicated in neuropathic pain and nicotine addiction^25–27^. Because *Gpr151* is a single-exon protein-coding gene, it was selected to explore the mRNA function by examining a single form of transcript, avoiding a need to consider alternatively spliced isoforms. We found that the 5’ untranslated region (5’UTR) of *Gpr151* mRNA promotes axon regeneration and identified RBPs interacting with the 5’UTR, including CSDE1, which was found to be a negative regulator of axonal regrowth in vitro. Moreover, we found that expressing an engineered form of the 5’UTR showed improved CSDE1- association and enhanced axon regeneration both in vitro and in vivo. Overexpression of the engineered 5’UTR did not significantly change the transcriptomic profiles but instead modulated the pool of CSDE1-associated RNA and promoted the release of RNA from CSDE1, including ribosomal protein-encoding mRNA. We also retrospectively re-analyzed our previous transcriptomic data to provide the ribosome-association efficiency of DLK-dependent DEGs that were previously published^28^. In this study, we introduce a new approach to injury-responsive DEG analysis by incorporating ribosome-association efficiency data from an axon regeneration paradigm and uncover the non-coding function of protein-coding mRNA as a biological modulator of the RBP function.

## Results

### *Gpr151* is an injury-responsive protein-coding gene that is not directed to ribosome

To identify injury-responsive genes (IRGs) regulating regeneration through their non-coding functions, *Advillin*-Cre mice crossed with *RiboTag* mice were utilized to specifically isolate the ribosome complex-associated RNA from the L4,5 dorsal root ganglia (DRG) with or without sciatic nerve crush injury (Fig. 1a)^8, 24, 29^. *Advillin*-Cre is an useful Cre mouse line expressing Cre recombinase from all subpopulations of peripheral sensory neurons without affecting other tissues in the organisms (Supplementary Fig. 1)^29^. In addition, *RiboTag* is an essential mouse line to isolate cell-type-specific ribosome as its cell expresses the hemagglutinin (HA) epitope-tagged ribosomal protein RPL22HA only when the cell express Cre recombinase^24^ (Fig. 1b). By using *RiboTag* x *Advillin*-Cre mice, we could isolate ribosome-associated mRNA from mice that had sciatic nerve crush injury (Injured) not (Uninjured) (Fig. 1c). Comparative analysis between the whole transcriptome (bulk-seq) and the ribosome-bound RNA (*RiboTag*-seq) showed that a large portion of upregulated coding genes was not directed to ribosome complex, suggesting that non-coding functions of these need to be investigated (Fig. 1d). Total 1,403 protein-coding genes identified from bulk-seq as the injury-responsive DEGs with statistical significance from edgeR analysis (*p*-value<0.05) and plotted with *RiboTag*-seq result. We found that the positive correlation of 62% (36%+26%) genes and the negative correlation of 38% (15%+23%) genes from the total 1,403 protein-coding injury-responsive DEGs (Supplementary Data 1).

**Fig. 1.**
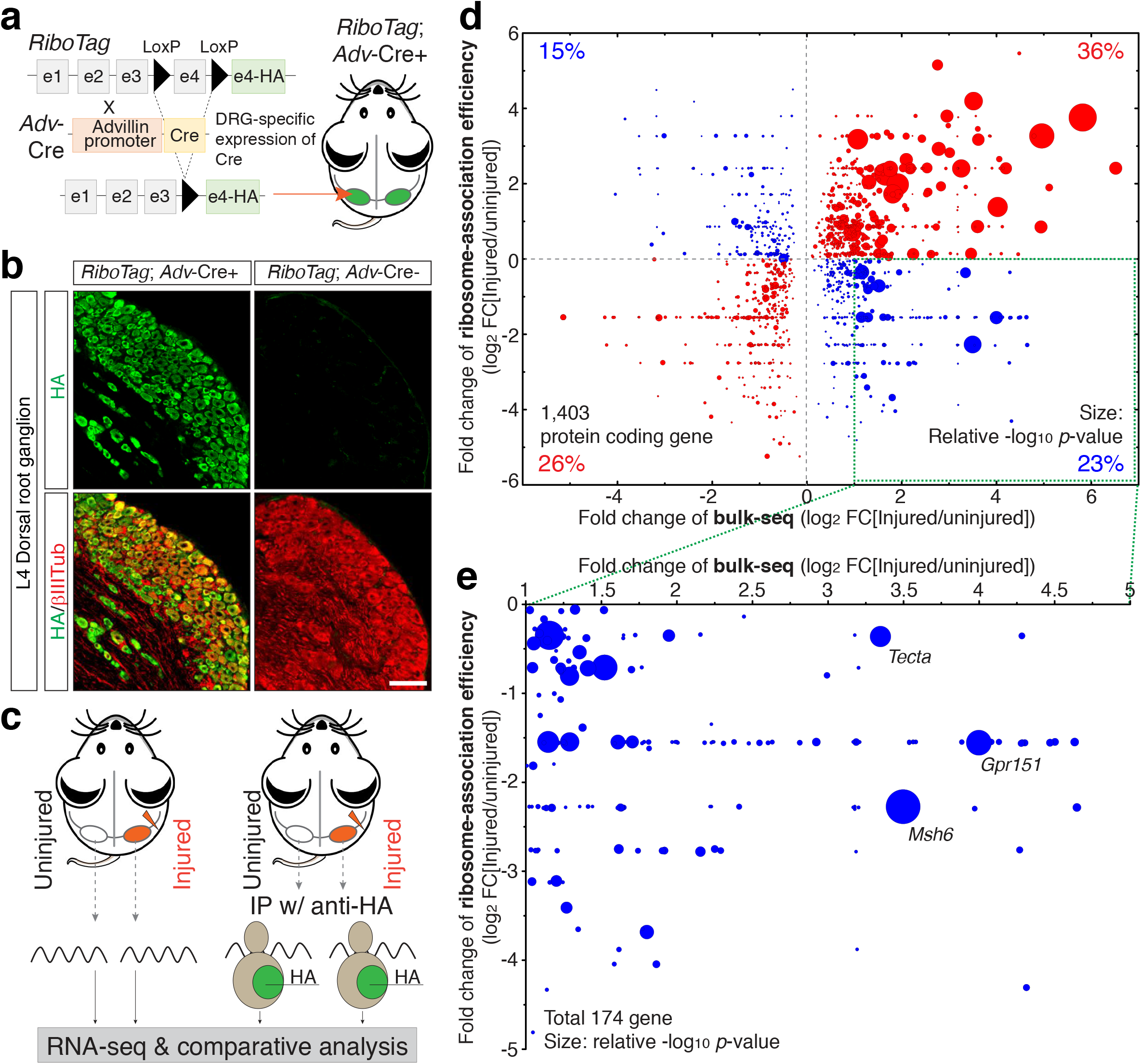
Comparative analysis between differential expression and ribosome-association revealed the distinct efficiency of ribosome-association of injury-responsive genes. (a) Schematic of the mouse genetics. *RiboTag* mouse carries the exon 4 of *Rpl22* gene (e4) that is floxed by *LoxP*. *Advillin*-Cre (*Adv*-Cre) line is a mouse line expressing Cre recombinase driven by *Advillin* promoter only from all subtypes of peripheral neurons. (b) Immunohistochemistry of mouse DRG sections stained with anti-HA and anti-βIII tubulin (Tub) antibody. *Advillin-*Cre mouse crossed with *RiboTag* mouse expresses HA-epitope tagged RPL22 specifically in peripheral sensory neurons. Mouse L4 DRG was dissected and its cryosection was immunostained with anti-HA (green) and βIII tubulin (red) antibodies. Scale, 100 μm. (c) Illustration of experimental scheme. Comparative analysis between differential gene expression and ribosome-association efficiency. IP, immunoprecipitation; HA, HA-epitope tag. (d) Fold change (FC) of bulk-seq, differential gene expression (x-axis, [Injured/uninjured]) was plotted against fold changes of differential ribosome-association efficiency (*RiboTag*-seq) (y-axis, [Injured/uninjured]) at 24 hours after sciatic nerve crush injury. The total 12,967 of protein-coding genes were identified and 1,403 DEGs with statistical significance (*p*-value < 0.05) were plotted. The size of circle indicated the relative -log_10_[*p*-value of FC of bulk-seq]. Red and blue color indicated the positive and negative co-relation of the fold changes between bulk-seq and *RiboTag*-seq results. The percentile indicated the counts of the genes with of bulk-seq fold changes *p*-value<0.05. (e) The genes with log_2_FC>2 from the quadrant 4 of (d) (total 174 gens) were re-plotted with x-axis of log_2_FC bulk-seq and y-axis of log_2_FC *RiboTag*-seq. *EdgeR*^36, 37^-R package was used for data processing to determine the statistical significance.

Among the genes that were highly upregulated over the two-fold with the negative efficiency of ribosome-association, we identified *Gpr151* as a potential candidate, an orphan G protein-coupled receptor that mediates nicotine sensitivity and neuropathic pain^25–27, 30, 31^ as it satisfied the important two factors. First of all, *Gpr151* displayed the high fold changes with high confidence after sciatic nerve injury as well as the absolute expression levels of *Gpr151* (logCPM and LR from *edgeR*) was reasonably high enough so possible to consider it works as RNA itself (Fig. 1e and Supplementary Fig. 2). Second, *Gpr151* mRNA displayed no significant changes in ribosome-association as its log_2_ FC_*RiboTag* was -1.6 with no statistical significance of *p*-value 0.4, even when its mRNA showed the dramatic upregulation (Fig. 1e and Supplementary Data 1).

While *Gpr151* upregulation after nerve injury has been reported by several transcriptome studies using DRG tissues, its role in the regulation of axon regeneration remains unknown ^27, 32–35^. The comparative analysis of bulk-seq with RiboTag-seq reveled that *Gpr151* mRNA was a potential candidate functioning as RNA independently as its protein product.

To validate the analysis, the expression levels of Gpr151 mRNA and its protein were investigated from mouse L4,5 DRG tissue lysates prepared at 24 hours with (+Injury) or without (-Injury) sciatic nerve crush injury. We found that western blot analysis, qPCR analysis, fluorescence in situ hybridization (FISH) and immunohistochemistry (IHC) data consistently supported the findings obtained by RNA-sequencing and bioinformatic analysis. While nerve injury greatly induced the *Gpr151* mRNA levels by RT-qPCR analysis (Fig. 2b) and FISH analysis (Fig. 2c), its protein levels were not upregulated by nerve injury but instead downregulated significantly as shown in western blot analysis (Fig. 2a) and IHC analysis (Fig. 2d). These results showed that *Gpr151* was one of the most upregulated injury-responsive gene that was not associated with ribosome complexes, suggesting that *Gpr151* was suitable for studying the non- coding function of a coding IRG for regulating axon regeneration. In addition to its injury responses, *Gpr151* was a single-exon gene and therefore it allowed us to trace a single transcript form without considering alternatively spliced isoforms.

**Fig. 2.**
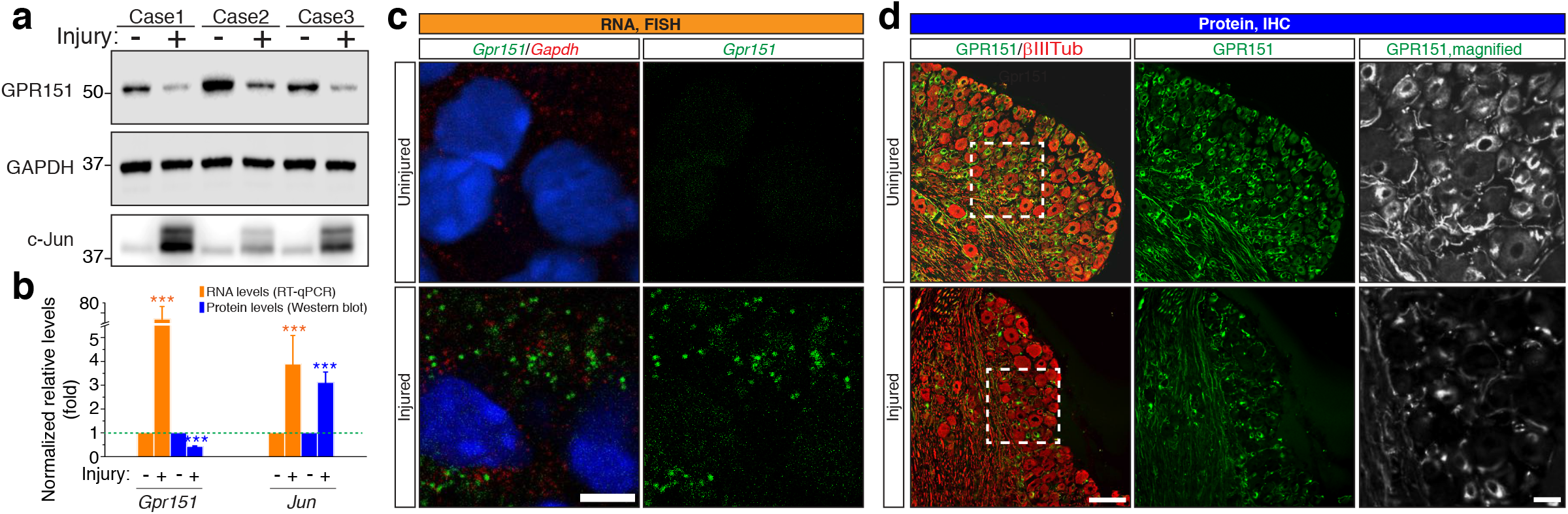
*Gpr151* mRNA was highly upregulated while its protein level was reduced. (a) Western blot analysis from mouse L4,5 DRGs tissue lysates with anti-GPR151, anti-GAPDH and anti-c-Jun antibody. L4,5 DRGs were dissected at 24 hours with (injury: +) or without (injury: -) sciatic nerve crush injury. (b) Normalized relative RNA or protein levels of *Gpr151*. *Jun* was used as an internal control to validate the sciatic nerve injury responses. Mouse L4,5 DRGs were dissected at 24 hours with (+) or without (-) sciatic nerve crush injury and subjected to analysis. Relative RNA levels were determined by RT-qPCR analysis (ÄÄCt) and normalized to the levels of *Gapdh* expression (orange box). Relative protein levels were determined by *ImageStudio* (Leica) and normalized to *Gapdh* levels (blue box). (n=3; mean±SEM; \*\*\**p*<0.001 by *t*-test). (c) Fluorescence in situ hybridization (FISH) analysis of L4 DRG sections with probes to *Gpr151* and *Gapd*h. DAPI solution was applied to visualize nucleus. Scale, 5 μm. (d) Immunohistochemistry (IHC) of L4 DRG with (injured) or without (uninjured) sciatic nerve crush was dissected, sectioned and immunostained with anti-GPR151 antibody (green) and anti-βIII tubulin (Tub) antibody (red). Scale bar, 100 μm.

### GPR151 protein-null mice showed enhanced axon regeneration after sciatic nerve injury

To investigate the role of *Gpr151* mRNA and its protein separately, we employed *Gpr151*-targeted mutant mice (KO) in which the *Gpr151* CDS was disrupted by insertion of a *LacZ* cassette to abolish GPR151 protein production. Although the KO mice did not express GPR151 protein, the engineered *Gpr151* gene had the part of *Gpr151* mRNA including both 5’UTR and 3’UTR (Fig. 3a). Therefore, the KO mouse line enabled to test the requirement of GPR151 protein for efficient axon regeneration. First, we tested the expression levels of the mutant *Gpr151* mRNA from KO mice with or without sciatic nerve injury. Total RNA was extracted from L4,5 DRGs dissected at 3 days after introducing injury by crushing the sciatic nerves with forceps as described in methods and subjected to RT-qPCR analysis (Figs. 3b, 3c).

**Fig. 3.**
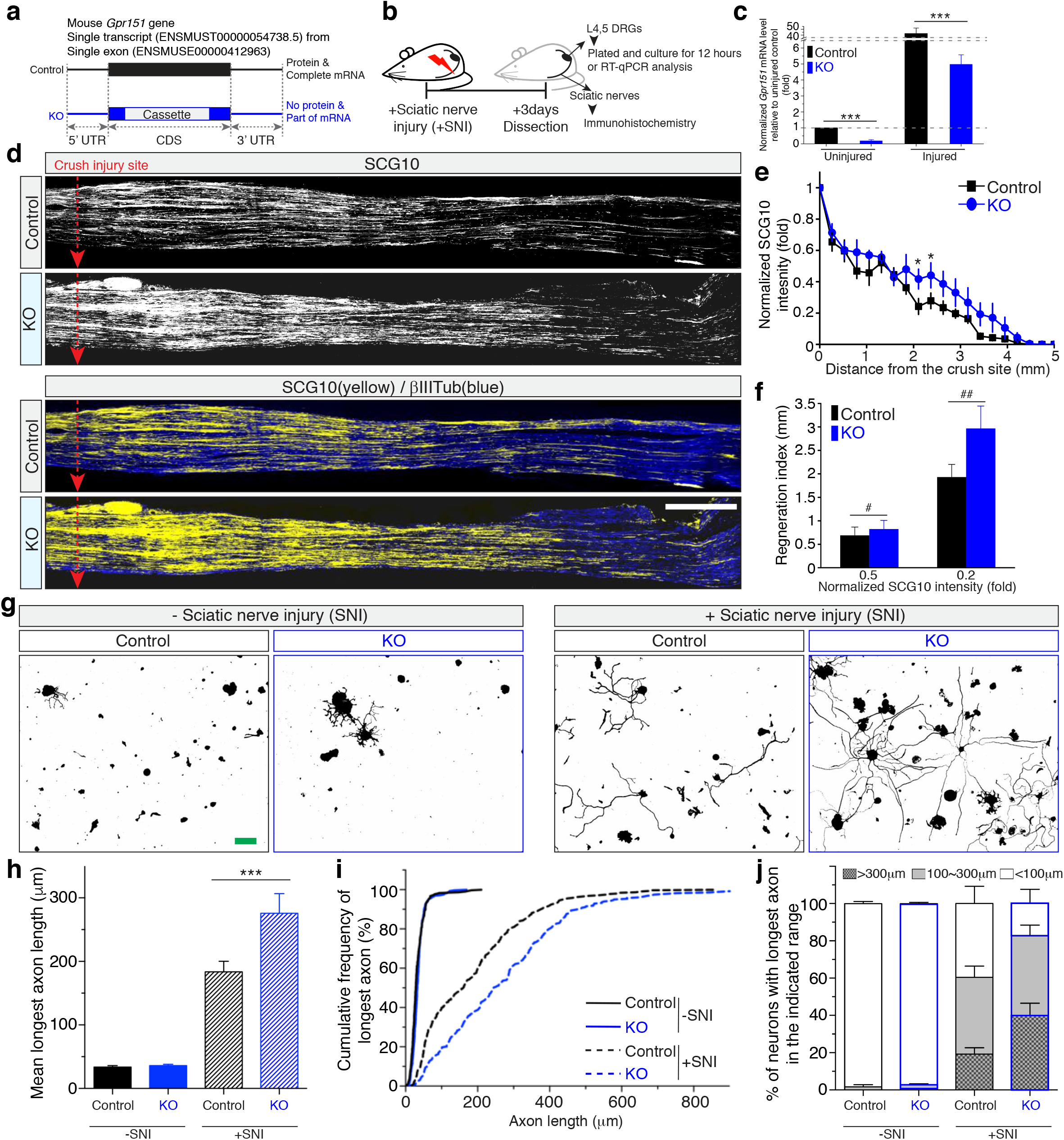
GPR151 protein-null mice showed enhanced axon regeneration after sciatic nerve injury. (a) Schematic structure of *Gpr151* gene in wild type control (black) or targeted knockout (KO) mice (blue). *Gpr151* gene was disrupted by the insertion of *LacZ*-cassette in the CDS region. (b) Schematic diagram of timeline illustrating the pre-conditioning injury and sample preparation. Sciatic nerves of wild type control or KO mice were injured by crushing the nerves with forceps and dissected at 3 days after injury. The sciatic nerves were prepared as cryosections and subjected to immunohistochemistry with anti-SCG10 and βIII tubulin antibodies. L4,5 DRGs were subjected to RT-qPCR analysis (ÄÄCt) or adult DRG neuron cultures. (c) RT- qPCR analysis of mouse L4,5 DRGs. Average of normalized levels of wild type control and KO mRNA levels at 3 days after introducing sciatic nerve crush injury. The averages of relative *Gpr151* mRNA levels from uninjured KO, injured control and KO samples was compared to the average level from control uninjured samples (set to 1). (n=6 for each condition; \*\*\**p*<0.001, ANOVA followed by *Tukey* test). (d) Representative longitudinal sections of immunohistochemistry of sciatic nerves from in vivo axon regeneration assay. The longitudinal cryosections were immunostained with anti-SCG10 (white; top and yellow converted; bottom), a marker protein specifically labelling regenerating axon. βIII tubulin antibody was used for counterstaining to visualize the nerve sections (blue; bottom). The red dotted arrow line indicates the crush site. Scale bar, 500 μm. (e) Average of normalized SCG10 intensity from immunostained sections in (d). Immunostained SCG10 fluorescence intensity was acquired by measurement windows with 100 pixel-width (equivalent to 87.7 μm) from crush sites to distal parts of nerves sections at an every 100-pixel distance from *ImageJ*. The acquired intensity from every measurement window was normalized to the intensity of the crush site (n=7 for control mice and n=5 for KO mice; \**p*<0.05 by *t*-test; mean ± SEM). (f) Regeneration index calculated from (e). Regeneration index was defined as a distance of an indicated SCG10 intensity. Average distance of normalized SCG10 intensity 0.5 or 0.2 was calculated from (e) and presented as regeneration index (mm). (^#^*p* = 0.59, ^##^*p* = 0.05 by *t*-test). (g) Mouse L4,5 DRG tissues were dissected at 3 days with (+SNI) or without (-SNI) sciatic nerve crush injury (SNI). DRG neurons were plated and cultured to monitor pre-conditioning effect-induced neurite outgrowth. Scale bar, 100 μm. (h) Average of the longest axon length from (g) (three independent biological replicates; total 6 mice; total 304, 313, 291, 283 cells for each condition, -SNI Control, -SNI KO, +SNI Control, +SNI KO; \*\*\**p*<0.001 by ANOVA followed by Tukey tests). (i) Cumulative frequency of the longest axon length from (g). (j) Percentage of neurons in three categories of the longest axon length.

When mice did not have the sciatic nerve crush injury, the basal level of *Gpr151* mRNA from KO DRGs was only 19.3% of that in wild type DRGs (Fig. 3c). We found the highly upregulated expression of *Gpr151* mRNA from L4,5 DRGs of wild type control mice by introducing sciatic nerve crush injury, showing an average of 40-fold change from wild type control mice. However, the induction from KO mice was 26-fold increase, which was much less than the change from the control mice (Fig. 3c). This result indicated that KO mice also upregulated the expression of the mutant *Gpr151* mRNA from L4,5 DRGs in response to sciatic nerve injury, although the expression level of mutant mRNA was much lower in KO mice comparing to wild type control mice.

Next, the regenerative capacity was monitored by in vivo axon regeneration assays using anti-SCG10 antibody^38, 39^, showing that the KO mice exhibited enhanced axon regeneration after sciatic nerve crush injury (Figs. 3d, 3e, 3f). Crush injury was introduced to sciatic nerves and dissected at 3 days after introducing injury for immunohistochemistry analysis. The longitudinal cryosections of crushed sciatic nerves from control and KO mice were stained with anti-SCG10 antibody that was specifically labeling regenerating axons and not axotomized distal part of injured axons ^38, 39^. Normalized intensity of SCG10 staining showed that the sections prepared from KO mice displayed higher levels of SCG10 immunofluorescence, indicating the more numbers of regenerating axons in crushed sciatic nerves (Fig. 3e). The distances showing the 0.5- and 0.2-fold of SCG10 intensity from the crush site were calculated from (e) and presented as the regeneration index (Fig. 3f), indicating GPR151 protein-null KO mice had no impairment of axon regeneration. Rather, KO mice displayed the mild enhanced axon regeneration after sciatic nerve crush injury. GPR151 protein-null mice did not show a significant change of the number of DRG neurons (Supplementary Fig. 3).

To monitor the neurite growth efficiency of the adult DRG neurons in vitro, we tested the pre-conditioning effect for neurite outgrowth from control and KO mice. L4,5 DRG tissues were dissected from control or KO mice at 3 days after sciatic nerve crush injury and plated on coated culture dishes for 12 hours in vitro. With no pre-conditioning potentiation (-SNI), the DRG neurons cultured from both the control and KO mice did not project their neurites efficiently (Fig. 3g, left). However, plated adult DRG neurons that had pre-conditioning injury (+SNI) extended their neurites extensively (Fig. 3g, right). Notably, DRG neurons from KO mice projected much longer neurites than those from the control mice (Figs. 3g, 3h, 3i). The number of DRG neurons of KO mice with neurite lengths shorter than 100 μm was less than 50% of those of the control mice, while the KO mice had two-fold more neurons with neurites longer than 300 μm compared to the control (Fig. 3j). These results showed that GPR151 protein was not required for axon regeneration in vivo and neurites outgrowth in vitro. Rather, GPR151 protein might have a negative function for axonal growth because GPR151 protein-null DRG neurons regenerated their axons in vivo more efficiently and extensively projected longer neurites in vitro.

### 5’UTR of *Gpr151* mRNA was required for efficient neurite outgrowth

Because GPR151 protein-null mice showed no impairment of axon regeneration, the requirement of *Gpr151* mRNA was tested from cultured embryonic DRG neurons by knocking down *Gpr151* mRNA using lentiviral transduction. To monitor neurite outgrowth efficiency, in vitro replating assay was done^40–44^ and revealed that *Gpr151* knockdown (KD) significantly reduced the potential of neurite extension when lentivirus-transduced neurons at DIV2 were detached and replated on new culture at DIV5 (Figs. 4a, 4c, 4f). Next, we tested if the expression of *Gpr151* mRNA or its protein restored the capacity of neurite outgrowth from *Gpr151* KD neurons using the artificial *Gpr151*-related lentiviral constructs CDSwt and CDSmut (fig. 4b). Both 5’UTR-CDSwt and -CDSmut lentiviral constructs expressed 5’UTR-CDS of *Gpr151* mRNA. However, the construct CDSmut had the point mutation at the original start codon AUG of CDSwt substituted to AUC to reduce its translation efficiency (Fig. 4d)^45–47^. Therefore, CDSmut expression recovered the expression of *Gpr151* mRNA with its 5’UTR and CDS parts with the minimal changes without overexpression of its protein product (Fig. 4e). To validate the experimental design, the overexpression levels of their mRNAs were tested by RT-qPCR analysis, showing that embryonic DRG neurons transduced by CDS, 5’UTR-CDSwt or 5’UTR-CDSmut lentivirus produced the similar levels of their mRNAs. In addition, these mRNA products were resistant from the degradation mediated by *Gpr151* shRNA that were designed to target the 3’UTR of *Gpr151* mRNA (Fig. 4j).

**Fig. 4.**
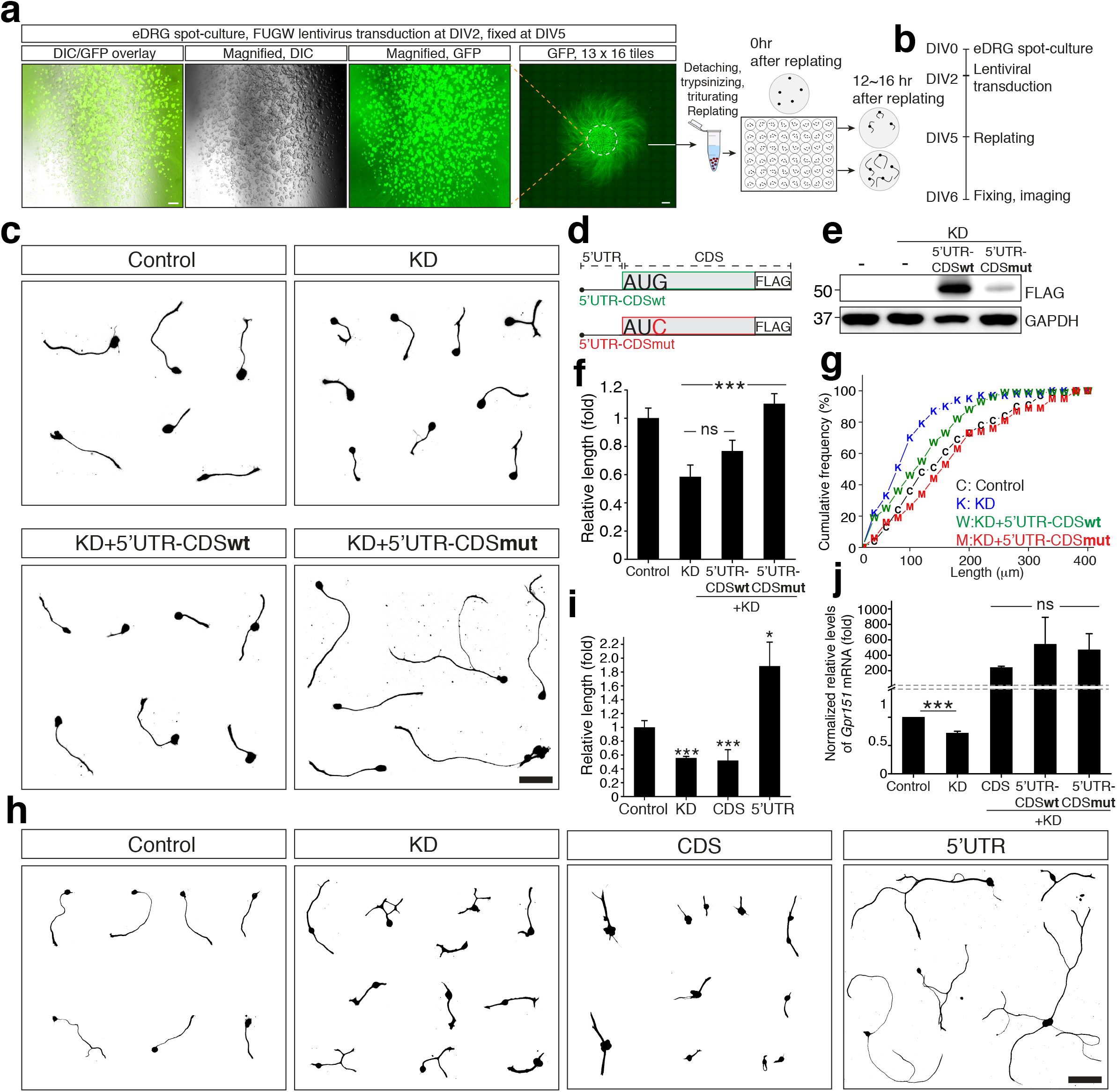
*Gpr151* mRNA was required for neurite outgrowth, while its protein inhibited neurite extension. (a) Illustration of embryonic DRG neurons spot-culture and in vitro replating assay. Mouse DRGs were dissected from E13.5 embryo and cultured as spot as described in methods. Spot-culture DRG neurons were transduced by GFP-expressing control lentivirus (FUGW) at DIV2 and fluorescence images were acquired at DIV5. Scale bar, magnified, 200 μm left, titled 500 μm right. To visualize the entire view of spot-culture neurons with their neurite projections, total 13 x 16 titled images were acquired and re-constructed as the single panel by stitching. GFP and differential interference contrast (DIC) images were acquired to show high efficiency of gene expression by lentiviral transduction. (b) Experimental timeline of in vitro gene delivery and replating assay. (c) In vitro replating assay of embryonic DRG neurons. All the in vitro gene deliveries were done by lentiviral transduction from spot-culture DRG neurons at DIV2 to achieve high efficiency of gene expression. Transduced neurons were then replated at DIV5 to monitor the change of neurite outgrowth efficiency from manipulated DRG neurons. Control (pLKO-scrambled); KD, *Gpr151* knockdown (pLKO-sh*Gpr151*); KD+5’UTR-CDSwt, *Gpr151*- knockdown with 5’UTR-CDSwt overexpression (pLKO-sh*Gpr151*+FUGW-5’UTR-CDSwt); KD+5’UTR-CDSmut, *Gpr151*-knockdown with 5’UTR-CDSmut overexpression (pLKO- sh*Gpr151*+FUGW-5’UTR-CDSmut). Scale bar, 100 μm. (d) Schematic diagram of the constructs 5’UTR-CDSwt and 5’UTR-CDSmut. “AUG” and “AUC” indicated the nucleotide sequences at the start codon position of GPR151*-*protein coding sequence (CDS). 5’UTR indicates 5’-untranslated region of mouse *Gpr151* gene (ENSMUSG00000042816). (e) Western blot analysis validating expression level of the constructs of 5’UTR-CDSwt and 5’UTR-CDSmut shown in (d). Anti-FLAG antibody was used for western blot analysis to specifically detect exogenous expressed GPR151 protein from (d). (f) Average of relative neurite length (total 67, 63, 67, 62 transduced cells for each condition from three biological replicates; \*\*\**p*<0.001 by ANOVA followed by Tukey tests; ns, not significant with *p*>0.05). (g) Cumulative frequency of (f). (h) In vitro replating assay of embryonic DRG neurons. Control (pLKO-scrambled+FUGW-null), *Gpr151* knockdown (KD, pLKO-sh*Gpr151*), coding sequence overexpression (CDS, FUGW-CDSwt) and 5’UTR of *Gpr151* overexpression (5’UTR, pLKO-5’UTR of *Gpr151*). Scale bar, 100 μm. (i) Average of relative neurite length (n=160, 147, 267, 161 cells for each condition; \**p*<0.05, \*\*\**p*<0.001 by ANOVA followed by Tukey tests). (c and h) The images were generated as collages by collecting individual neuron from acquired raw microscopic fields and placing the images altogether in a single blank panel. The representative individual cell images from each condition were subjected to analysis and collages by selecting a cell that was not overlapped from adjacent cells to measure neurite length. The original acquired raw microscopic fields were in Supplementary Fig. 5-12. (j) Normalized relative levels of *Gpr151* mRNA by RT-qPCR analysis (n=3; \**p*<0.05, \*\*\**p*<0.001 by *t*-test). Control, pLKO-scrambled; KD, pLKO-*shGpr151*; CDS+KD, FUGW- CDSwt+pLKO-sh*Gpr151*; 5’UTR-CDSwt+KD, FUGW-5’UTR-CDSwt+pLKO-sh*Gpr151*; 5’UTR-CDSmut+KD, FUGW-5’UTR-CDS+pLKO-sh*Gpr151*.

Next, in vitro replating assay was done from *Gpr151* knockdown neurons co-transduced by 5’UTR-CDSwt or 5’UTR-CDSmut lentivirus to monitor the efficiency of neurite extension. We found that 5’UTR-CDSwt-transduction failed to recover the original capacity of neurite extension from endogenous *Gpr151* knocked down (KD) neurons. In contrast, the lentiviral delivery of 5’UTR- CDSmut to KD neurons successfully restored the impaired neurite extension (Figs. 4c, 4f, 4g). These results indicated that *Gpr151* mRNA and its protein involved in neurite projection from different ways and overproduction of GPR151 protein did not promote neurite growth efficiency. This result implied that overexpressing *Gpr151* mRNA by constructs that also induced GPR151 protein overproduction failed to enhance neurite outgrowth, as we showed that GPR151 protein might function as a negative regulator from KO mice (Fig. 2). Rather, expressing *Gpr151* mRNA with no overproduction of its protein restored the neurite growth capacity from KD neurons (Figs. 4c, 4f).

As we found that *Gpr151* mRNA was required and responsible for the restoration of the efficiency of neurite growth from KD neurons, 5’UTR or CDS was overexpressed in embryonic DRG neurons to specify the part of mRNA involved in enhancement of neurite growth efficiency. In vitro replating assay showed that 5’UTR-lentivirus transduced neurons extensively projected longer neurites than those from control neurons (Figs. 4h, 4i). However, transduction of CDS-encoding lentivirus did not promote growth capacity. Rather, CDS-lentivirus transduction reduced the projection efficiency of neurites significantly as much as *Gpr151* knockdown did. To further validate, *Gpr151*- knockdown neurons were restored by co-transducing CDS-lentivirus or 5’UTR-lentivirus, showing that the original capacity of neurite growth was recovered only by 5’UTR expression while restoration of CDS expression failed to recover (Supplementary Figs. 4a, 4b). This observation indicated that the introducing 5’UTR of *Gpr151* mRNA was sufficient for promoting neurite outgrowth and *Gpr151*- invovled neurite outgrowth was mediated via its 5’UTR region. Additionally, overproducing GPR151 protein might be detrimental to neurite growth, implying that adult DRG neurons of mouse might have a mechanism to differentially regulate the injury-responsive production of *Gpr151* mRNA and GPR151 protein in the PNS such as DRG neurons^48^.

### The 5’UTR of *Gpr151* binds to CSDE1, a negative regulator of neurite extension

To understand the molecular mechanism of *Gpr151*-5’UTR-mediated neurite outgrowth, 5’UTR-interacting proteins were screened by pull-down assay followed by mass spectrophotometry analysis. A total of 25 proteins were identified with the high confidence scores and the matched molecular weights (Fig. 5a, Supplementary Data 2). Among them, CSDE1 was recognized as a potential candidate to investigate because of its known neuronal roles. CSDE1 is a negative regulator of neural differentiation in stem cells and loss of CSDE1 accelerates neurogenesis^49^. In addition, likely gene-disrupting variants in CSDE1 are associated with autism spectrum disorder, indicating that CSDE1 has the functional role in neurons^50^. Notably in this study, Guo et al. showed that CSDE1 knockdown promotes neurite and axon outgrowth^50^. These results stimulated us to hypothesize that CSDE1 was a key effector for *Gpr151* mRNA-mediated neurite outgrowth by functioning as a non-ribosomal RNA-binding protein (RBP) of *Gpr151* mRNA. Moreover, we found that CSDE1-binding motif was evolutionary conserved in mouse, rat and human *GPR151* 5’UTR region at a similar distance from the upstream of start codon, implying that CSDE1 was a potential mediator for *Gpr151* mRNA function^49, 51, 52^ (Fig. 5b).

**Fig. 5.**
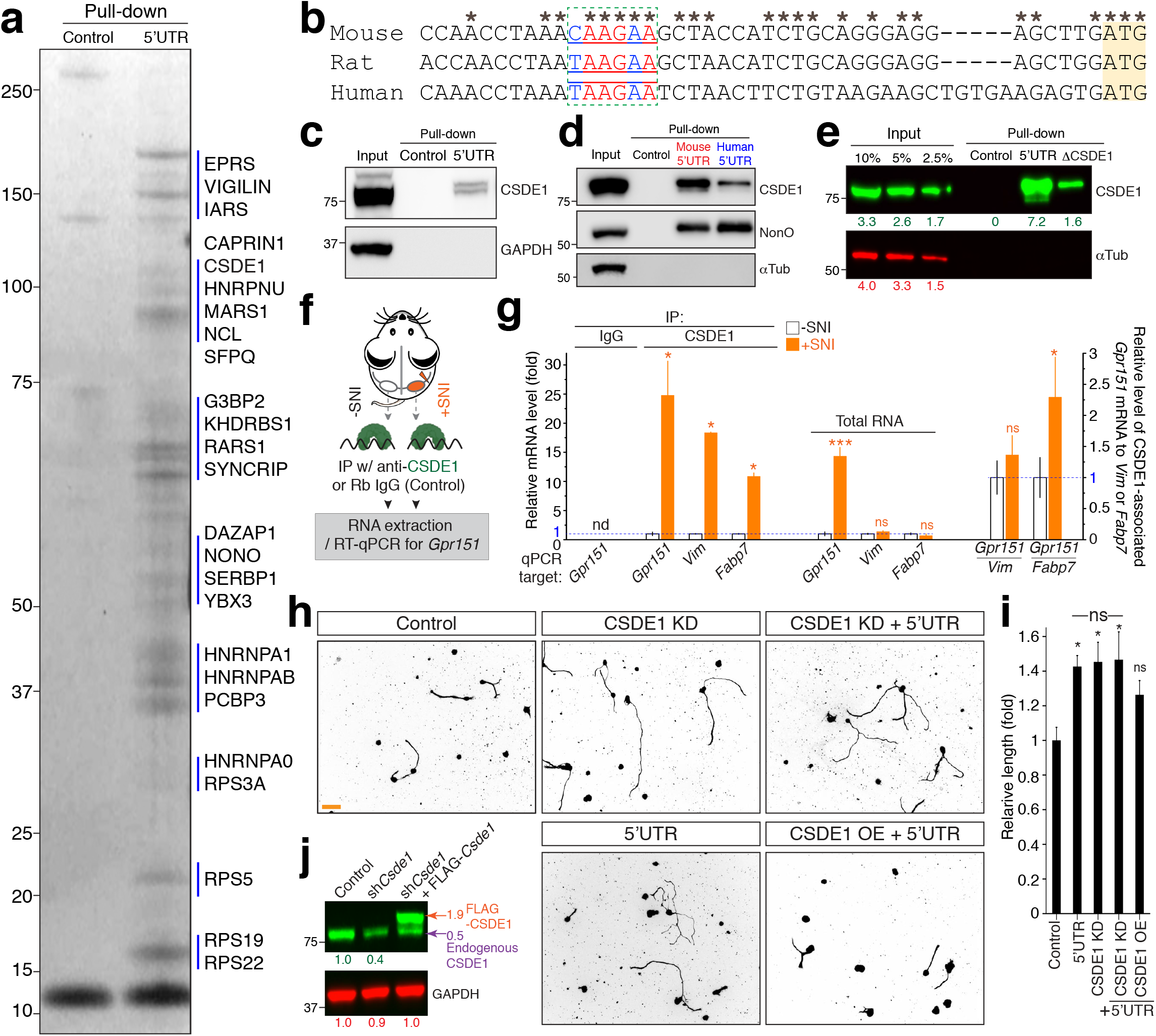
CSDE1 was a negative regulator of neurite outgrowth and mediated *Gpr151*-5’UTR- induced neurite extension. (a) Coomassie staining image of SDS-PAGE from pull-down assay. Control, Biotin-AMP; 5’UTR, Biotin-*Gpr151* 5’UTR RNA. The labels indicated the identified proteins from mass spectrophotometry analysis. The identified peptides and the scores were in Supplementary Data 2. (b) Alignment of 5’UTR sequences of mouse, rat and human *GPR151* gene. Green box, nAAGnA, CSDE1-binding consensus motif^49, 53^, yellow shadow, the start codon. (c) Western blot analysis of pull-down assay. Total protein lysates prepared from embryonic cultured DRG neurons were incubated with control (Biotin-AMP) or 5’UTR (Biotin-5’UTR-41mer RNA) for pull-down assay and subjected to western blot analysis with anti-CSDE1 and anti-GAPDH antibody. (d) Western blot analysis of pull-down assay with the baits, mouse 5’UTR or human 5’UTR RNA of *GPR151*. NonO, a nuclear RNA-binding protein, one of the identified binding proteins was also immunoblotted as an internal control. (e) Western blot analysis of pull- down assay with the baits, mouse 5’UTR or ΔCSDE1 RNA. ΔCSDE1 RNA was the CAAGAA- CSDE1-binding motif -deleted RNA. The numbers indicated the fluorescence intensities of the corresponding immunoblot bands from the infrared dye (IR-dye) detection with *Odyssey* (Leica) and measured by *ImageStudio* software (Leica). (f) Illustration of CSDE1 immunoprecipitation and RT-qPCR analysis for (g). Mouse sciatic nerves injury was introduced by crushing and L4,5 DRGs were dissected at 24 hours after injury. Total tissues extracts were prepared and incubated with anti-CSDE1 antibody or normal rabbit IgG isotype control. The immunoprecipitants were subjected to RNA extraction and RT-qPCR to analyze the relative enrichments of *Gpr151* mRNA- associated with CSDE1 from uninjured or injured DRG tissues. The RNA-seq analysis results were indicated as “Total RNA” along the IP result with injury-dependent fold change (Fig. 1). (g) Relative enrichment of *Gpr151* mRNA levels analyzed by RT-qPCR (n=3; \*\*\**p*<0.001 by t-test; nd, not detected; mean±SEM). IP, immunoprecipitation; IgG, normal rabbit IgG isotype control antibody; black, uninjured, gray. injured. *Vim* and *Fabp7* were the internal controls genes known to interact with CSDE1^49^, showing that they did not respond to sciatic nerve injury. (h) In vitro replating assay of control (scrambled), 5’UTR overexpressed (5’UTR), CSDE1 knocked down (CSDE1 KD), 5’UTR and CSDE1 overexpressed (5’UTR+CSDE1 OE), 5’UTR overexpressed and CSDE1 knocked down (5’UTR+CSDE1 KD) neurons. The images were representative images of raw microscopic fields. Scale bar, 100μm. (i) Average of relative neurite length of (h). (n=4 replicates; total cell numbers analyzed; n=133, 132, 133, 131, 115 for control, 5’UTR, CSDE1 KD, 5’UTR+CSDE1 KD, 5’UTR+CSDE1 OE; \**p*<0.05, \*\**p*<0.01, ns, not significant by *t*-test; mean±SEM). (j) Western blot analysis of CSDE1 from cultured embryonic DRG neurons lysates. Lentivirus from pLKO-sh*CSDE1* and/or FUGW-CSDE1-myc/DDK was transduced to the cultures at DIV2 to knock down and/or overexpress CSDE1. The numbers indicated the fluorescence intensities of the IR-dye western blot bands. Endogenous and exogenous CSDE1 was labeled and indicated as arrows.

First, the interaction between the 5’UTR and CSDE1 was confirmed by RNA pull-down assay using biotinylated 5’UTR as a bait, followed by western blot analysis using an anti-CSDE1 antibody (Fig. 5c). As the binding motif was conserved in human 5’UTR *GPR151* mRNA, its interaction with CSDE1 was also validated by RNA pull-down assay using biotin-5’UTR RNA of human *GPR151* (Fig. 5d). NonO, a nuclear RNA-binding protein, another 5’UTR-binding protein identified from the screen, was also immunoblotted from the pull-down analysis to validate the mass spectrophotometry result (Fig. 5d). To test the requirement of the binding motif for the interaction, ⊗CSDE1, the binding motif-deleted mutant RNA was subjected to RNA pull-down assay. Although the mutant still interacted with CSDE1, deletion of the binding motif from 5’UTR significantly reduced the binding efficiency down to 0.2-fold of the control level (7.2 to 1.6), suggesting that this motif contributed to the stable association with CSDE1 (Fig. 5e).

To further validate the interaction in vivo, the association of endogenous *Gpr151* mRNA and CSDE1 protein was reversely verified by in vivo immunoprecipitation analysis from mouse adult DRG tissues. CSDE1 was immunoprecipitated from mouse L4,5 DRG tissues dissected at 24 hours with (+SNI) or without (-SNI) sciatic nerve crush injury and its immunoprecipitants were subjected to RT-qPCR analysis to quantitate CSDE1-associating *Gpr151* mRNA (Fig. 5f). We found that a dramatic increase of *Gpr151* mRNA from the CSDE1-immunoprecipitants prepared from mouse L4,5 DRG tissues with sciatic nerve injury (+SNI) (Fig. 5g). The sharp increase in the CSDE1-associated *Gpr151* mRNA showed a clear contrast to its affinity to ribosome complexes, which was unchanged after injury (Figs. 1d, 1e). From the study dissecting the role of CSDE1 in neuronal differentiation, *Vim* or *Fabp7* mRNA was identified as the major targets of CSDE1 regulation and associated with CSDE1^49^. Therefore, we tested the level of CSDE1-associated *Vim* or *Fabp7* mRNA, showing that both mRNAs were detected from the CSDE1- immunoprecipitants and they were enriched from the DRG tissues prepared with SNI. Notably, the levels of both mRNAs were not upregulated by SNI in vivo in contrast to *Gpr151*, indicating that *Gpr151* mRNA is a major regulator of CSDE1 in response to axonal injury selectively in vivo (Fig. 5g).

Although CSDE1 had the roles in neurons^49, 50^, there was no evidence showing that CSDE1 played a role in axon regeneration via interacting specific targets such as *Gpr151*. Therefore, we investigated the role of CSDE1 for *Gpr151* 5’UTR-mediated neurite outgrowth. FUGW-*Csde1*-myc/DDK lentivirus or pLKO-sh*Csde1* lentivirus was produced to overexpress CSDE1 protein or knock down *Csde1* in cultured embryonic DRG neurons, respectively. Using the lentivirus, we made a combinatorial analysis with manipulating CSDE1 expression. First, we tested the neurite outgrowth from *Csde1* knocked down neurons and found that *Csde1* knockdown (CSDE1 KD) promoted neurite outgrowth, showing the successfully replicating the previous result^50^. The enhancement level of neurite growth was comparable to the level from the 5’UTR-overexpressing neurons with no significant difference (Figs 5h, 5i, 4f, 4g). When 5’UTR was overexpressed in *Csde1* knockdown neurons, the neurons did not show synergistical effect to further enhance neurite outgrowth (Figs. 5h, 5i). In addition, 5’UTR-induced neurite outgrowth was counterbalanced by co-expression of CSDE1. This result suggested that CSDE1 had a potential function as a negative regulator of neurite outgrowth and implied that the 5’UTR of *Gpr151* interacted with CSDE1 to modulate its function in injured or regenerating axons. In addition, the result indicated that CSDE1 was the target RBP to understand the molecular mechanism in injured neurons.

### 5’UTR of *Gpr151* modulated CSDE1-associated RNA pool

As 5’UTR-induced neurite outgrowth was mediated by CSDE1, we asked whether CSDE1 was the major effector of *Gpr151*-5’UTR. To investigate the hypothesis, 5’UTR RNA sequences were engineered to improve the binding efficiency to CSDE1. The prediction of the secondary RNA structure of 5’UTR showed that the CSDE1-binding motif was partially double-stranded in the stem region (Fig. 6a). Because CSDE1 was known to bind to single-stranded RNA^53, 54^, we generated a mutant “5’UTRm” that was the full 5’UTR sequence but with the AGCUU substituted to its complementary sequence UCGAA so that the binding motif became single-stranded. In addition, “h”, the shortest 11-mer RNA with no CSDE1-binding motif and “⊗”, the 5’-part of the 5’UTR including the CSDE1-binding motif but no stem-loop structure was generated as controls (Fig. 6a).

**Figure 6.**
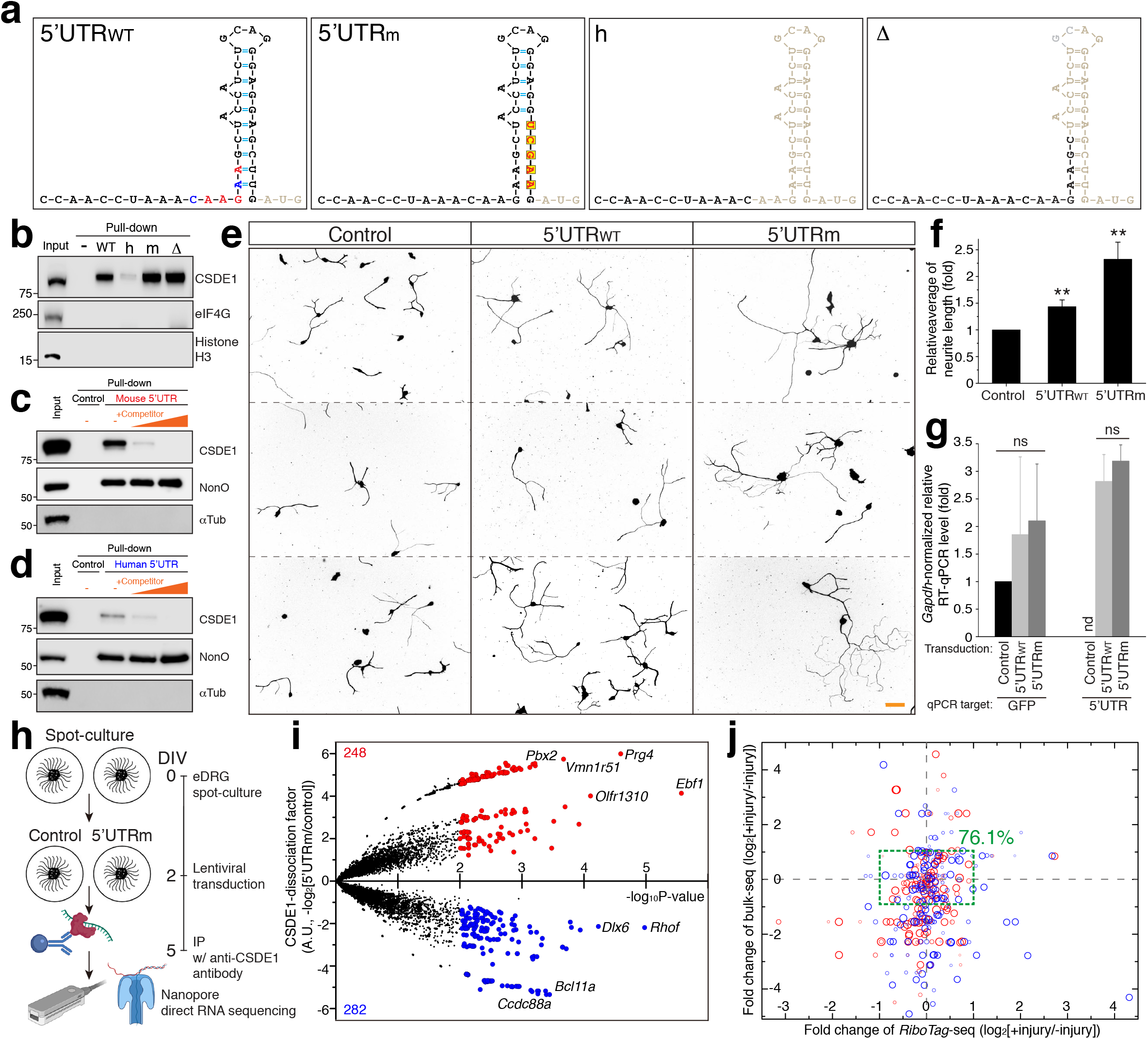
Expressing 5’UTRm of *Gpr151* promoted neurite outgrowth and modulated the CSDE1-associated RNA pool. (a) Illustration of the secondary structure of 5’UTR of *Gpr151* wild type (5’UTR_WT_), mutant 5’UTR with single-stranded CSDE1-binding motif (5’UTRm), no CSDE1-binding motif and no stem-loop (h), single-stranded and no stem-loop (Δ) mutant. (b) Western blot analysis of pull-down assay with the baits of (a). (c and d) Western blot analysis of pull-down assay with the competitor Δ for mouse and human 5’UTR. The competitor RNA Δ was synthetic RNA identical to (a) without biotin-conjugation. NonO was immunoblotted as an internal control of pull-down analysis. (e) Representative images of in vitro replating assay result of control, 5’UTR and 5’UTRm. Scale bar, 100μm. (f) Statistical analysis of relative average axon length from (e) (n=97, 104, 98 cells for control, 5’UTR_WT_ and 5’UTRm; \**p*<0.05, \*\*\**p*<0.001 by ANOVA followed by Tukey tests; mean±SEM). (g) *Gapdh*-normalized level of GFP, 5’UTR_WT_ and 5’UTRm by RT-qPCR analysis. GFP was expressed under CMV promoter from the second ORF in the lentiviral vector and its mRNA level was monitored as an internal control of viral transduction efficiency (n=3 replicates; ns, not significant by ANOVA followed by Tukey tests; mean±SEM). (h) Illustration of experimental time line of spot-culture, immunoprecipitation and Nanopore direct RNA sequencing. Mouse embryonic DRG neurons were cultured by spot-culture method as described. Spot-cultured neurons were infected by control or 5’UTRm lentivirus at DIV2 and lysed for CSDE1-immunoprecipitation at DIV5. RNA was extracted from the immunoprecipitants and subjected to library preparation for Nanopore direct RNA sequencing. (i) Volcano plot with the x-axis of -log_10_*P*-value and y-axis of CSDE1-dissociation factor. Two replicates of the sequencing results were subjected to *edgeR* analysis for analyzing differential level of transcripts associated with CSDE1. *P*-value was calculated from log_2_-fold change with two biological replicates. CSDE1-dissociation factor was defined as -log_2_[CSDE1-associated transcript level from 5’UTRm-expressing neurons / CSDE1-associated transcript level from control neurons]. Red dot indicated transcripts dissociated from CSDE1 when 5’UTRm was overexpressed (248 transcripts). Blue indicated transcripts showing enhanced association with CSDE1, reversely (282 transcripts). See also Supplementary Data 3. (j) The red and the blue transcripts from (i) was re-plotted with the x-axis of log_2_FC of *RiboTag*-seq result and the y-axis of log_2_FC of bulk-seq result. Green box indicated the transcripts with both values less than 1, 76.1% of total transcripts (248+282).

Pull-down assay showed that the bait 5’UTRm and Δ both made more stable association as the motif was single-stranded, while the bait h failed to bind to CSDE1, confirming the contribution of the motif sequence for the interaction (Fig. 5b). Therefore, RNA Δ with no biotin conjugation was used as a molecular competitor for pull-down competition assays to validate the interaction between 5’UTR with CSDE1, showing that addition of the competitor impaired the interaction for both mouse and human *GPR151* (Figs. 6c, 6d). Next, the efficiency of the neurite outgrowth of 5’UTR_WT_ and 5’UTRm-overexpressing neurons was monitored to know whether the enhanced CSDE1-binding affinity contributed to neurite outgrowth efficiency. In vitro replating assay result showed that 5’UTR_WT_ promoted neurite outgrowth by 150% to the control (Figs. 6e, 6f), consistent with the results of the initial assay (Figs. 4h, 4j). However, 5’UTRm expression showed more robust neurite extension promoted by 230% to the control (Figs. 6e, 6f). To verify that the enhanced function of 5’UTRm required the association with CSDE1, a CSDE1-binding motif-deleted mutant of 5’UTRm (5’UTRmÄCSDE1) was tested. We found that the mutant 5’UTRmÄCSDE1 failed to promote neurite outgrowth as much as 5’UTR_WT_ did (Supplementary Fig. 13), suggesting the function of 5’UTR was dependent on its physical association with CSDE1. The efficiency of lentiviral transduction and the expression level of 5’UTR_WT_ and 5’UTRm was analyzed by RT-qPCR assay, which showed that the levels were comparable with no significant difference, indicating that the differential neurite growth efficiency between 5’UTR_WT_ and 5’UTRm was not due to the different expression level (Fig. 6g).

To know the mechanism, we investigated CSDE1-associated RNA from control or 5’UTRm-overexpressing DRG neurons to compare (Fig. 6h). CSDE1 was immunoprecipitated from cultured embryonic DRG neurons and subjected to RNA extraction and Nanopore direct RNA sequencing analysis. The relative level of CSDE1-association was calculated and presented as CSDE1-dissociation factor of -log_2_[5’UTRm/control] (Fig. 6i). We identified 248 protein-coding transcripts that were dissociated from CSDE1 when 5’UTRm was overexpressed (Fig. 6i, red dots). Pathway analysis showed that these transcripts were included in Ras, Rap1 or FoxO signaling pathway (Supplementary Data 3). Interestingly, a 38% of the transcripts was membrane proteins, implying that membrane proteins were a potential group of targets regulated by the interaction between 5’UTR and CSDE1. In addition, we found that a group of transcripts showed increased efficiency for binding to CSDE1 when 5’UTRm was overexpressed (Fig. 6i). 282 transcripts were identified to be more recruited to CSDE1 by 5’UTRm overexpression. These transcripts were related to the functions of RNA transport, transcriptional regulation, mTOR pathway and apoptosis (Supplementary Data 3). This analysis showed that 5’UTRm overexpression induced a shift of the CSDE1-binding RNA pool. Notably, most of the 5’UTRm- regulated target mRNAs displayed no significant change nor from bulk-seq nor *RiboTag*-seq analysis in response to sciatic nerve injury (Fig. 6j). 76.1% of total identified transcripts showed less than two-fold change of both bulk-seq and *RiboTag*-seq analysis. This result implied that these transcripts might not be recognized from comparative injury-responsive DEG analysis and potential candidates for studying the neuronal responses to injury.

### Expression of the engineered 5’UTR RNA promoted axon regeneration in vivo

As we identified *Gpr151* as a regeneration associated genes (RAGs) functioning by 5’UTR of its mRNA, we overexpressed the engineered 5’UTR of *Gpr151* to test the efficiency of axon regeneration in vivo. The control scrambled 48-mer or the 5’UTRm 41-mer was packaged to adeno-associated virus (AAV, serotype 9) for in vivo gene delivery (Fig. 7a). For PNS axon regeneration, sciatic nerve was crushed to introduce injury at 12 weeks after virus injection and dissected at 3 days after injury for assessing axon regeneration efficiency (Fig. 7a). L4,5 DRGs were dissected and immunostained with βIII tubulin antibody to confirm the neuronal expression of GFP as an indicator of gene delivery (Fig. 7b). SCG10 antibody was used for immunohistochemistry to specifically label regenerating axons, which showed the robust axon regeneration in sciatic nerves from AAV-5’UTRm-injected mice (Fig. 7c). The normalized intensity of SCG10 fluorescence was much higher from AAV-5’UTRm mice than it from control mice at all the distances of the distal parts (Fig. 7d). Regeneration index was calculated as the distance showing 0.5 or 0.2-fold of SCG10-labeling intensity and showed that 5’UTRm-overexpression significantly promoted the efficiency of axon regeneration in sciatic nerves (Fig. 7e).

**Fig. 7.**
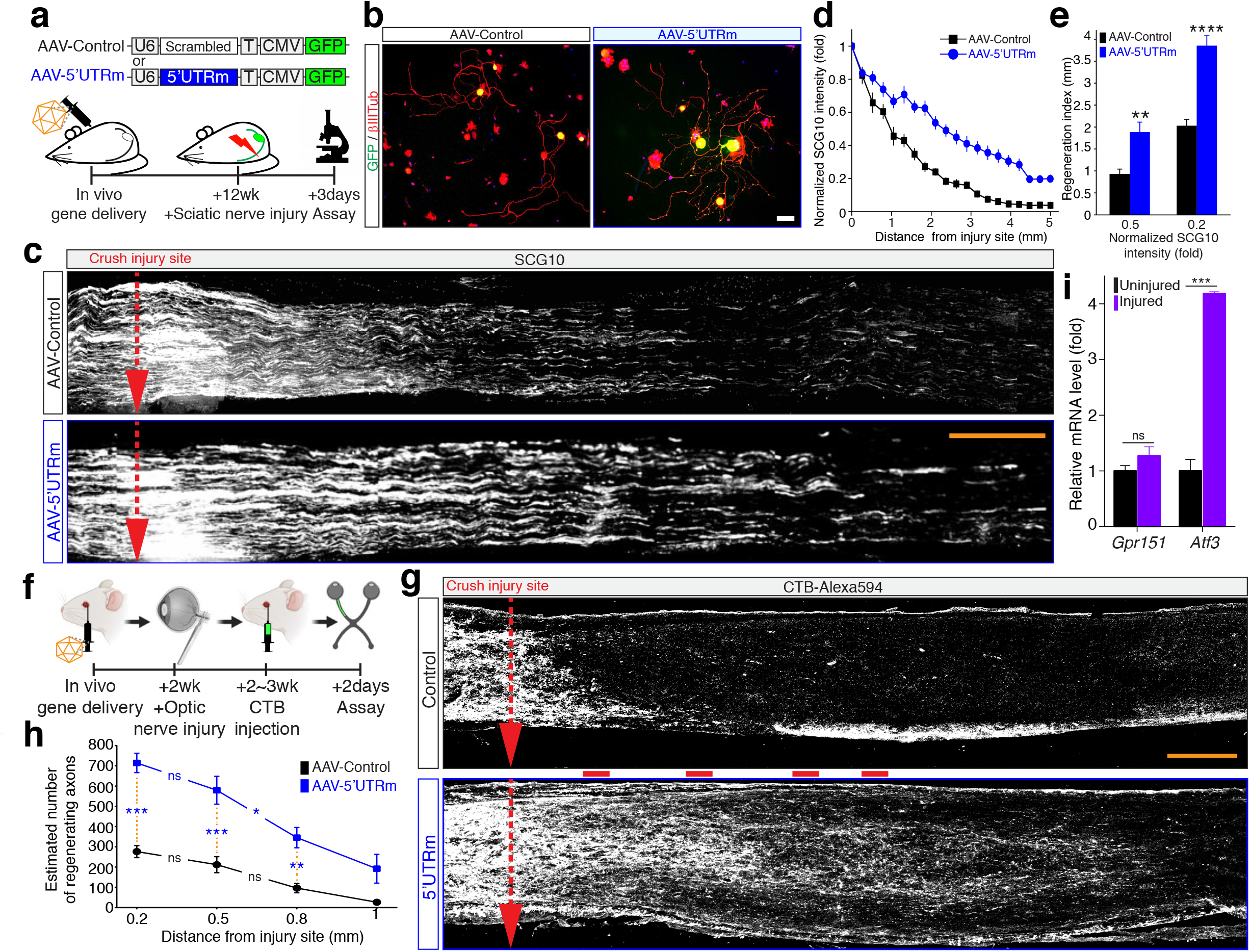
In vivo gene delivery of the engineered *Gpr151* 5’UTR RNA promotes axon regeneration. (a) Schematic diagram of AAV-control and AAV-5’UTRm adeno-associated virus vector and experimental timeline of in vivo axon regeneration assay in sciatic nerves (wk, weeks). U6, U6 promoter; Scrambled shRNA sequence 48-mer; T, transcription termination sequence; CMV, human cytomegalovirus immediate early enhancer/promoter; GFP, enhanced green fluorescence protein CDS. (b) Representative image of adult DRG neurons. L4,5 DRGs were dissected from AAV-injected mice and cultured for 12 hours and immunostained with βIII tubulin antibody. GFP fluorescence signal indicated AAV-mediated in vivo gene delivery and expression of target genes. Scale bar, 100 μm. (c) In vivo axon regeneration assay from crushed sciatic nerves. Representative longitudinal sections of sciatic nerves from AAV-control or AAV-5’UTRm- injected mice. Red dotted arrows indicate the injury site. Scale bar, 500μm. (d) Normalized SCG10 intensity measured from SCG10 immunostained sections in (c). Measurement window (100-pixel width, equivalent to 87.7 μm) was placed at the crush site and intensities were acquired at every 100-pixel to the distal part of the sections. (e) Regeneration index calculated from (d). (n=8 for AAV-control, 10 for AAV-5’UTRm; \*\**p*<0.01, \*\*\*\**p*<0.0001 by *t*-test; mean±SEM). (f) Experimental timeline of in vivo axon regeneration assay in optic nerves (wk, weeks). (g) Representative longitudinal sections of optic nerves from AAV-control or AAV-5’UTRm-injected mice. Regenerating axons were labeled by Alexa594-conjugated CTB injection. Red dotted arrows indicate the injury site. Red bars indicated the locations analyzed the estimation of regenerating axon numbers in (h). Scale bar, 200μm. (h) Estimated numbers of regenerating axons (n=5 for AAV-control, 6 for AAV-5’UTRm; \**p*<0.05, \*\**p*<0.01, \*\*\**p*<0.001 by one-way AVONA with Bonferroni test; mean±SEM). (i) Relative expression level of *Gpr151* and *Atf3* after injury in mouse retina tissues (injured / uninjured). RT-qPCR analysis of *Gpr151* and *Atf3* from mouse retina tissues dissected at 3 days with optic nerve crush injury (injured, purple) or without injury (uninjured, black) (n=3 for each condition; \*\*\**p*<0.001, ns, not significant by t-test; mean±SEM).

**Fig. 8.**
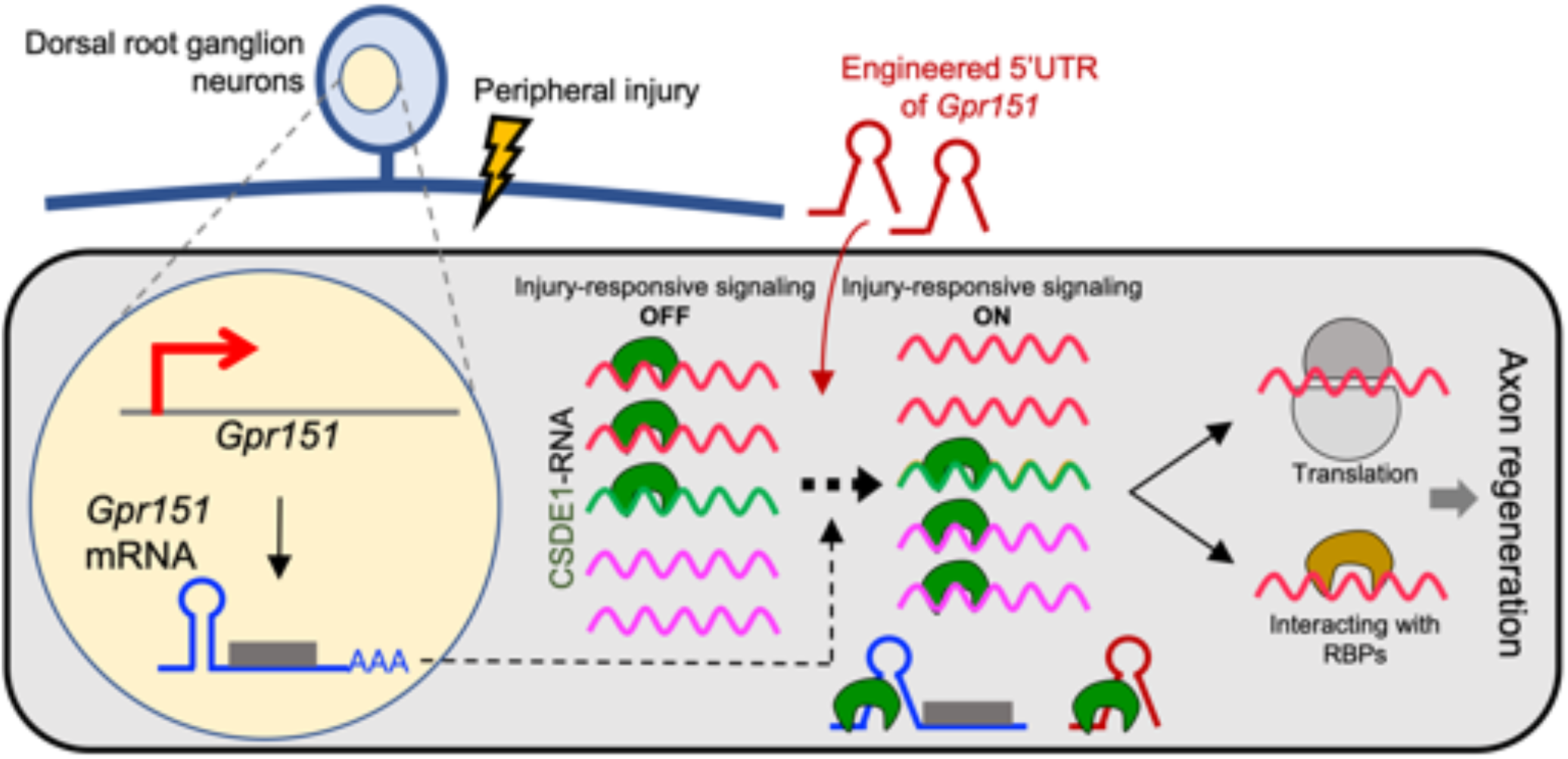
Illustration of the proposed model. *Gpr151* mRNA was upregulated in DRG neurons in response to sciatic nerve injury. Instead of directing to ribosome complexes for its translation, *Gpr151* mRNA interacted with RNA-binding proteins including CSDE1 via its 5’UTR and induced a change of the CSDE1-associated RNA pool. Driving this process by in vivo gene delivery of the engineered 5’UTR RNA amplified the regenerative potential on injured neurons from DRG tissues and retina ganglion cells.

Because neuronal responses to injury were differently regulated in the PNS and the CNS^1, 48, 55^, we tested if AAV-5’UTRm delivery was also effective in optic nerves for axon regeneration by optic nerve crush injury model (Fig. 7f) . Mouse optic nerve injury was introduced by crushing optic nerves with forceps at 2 weeks after AAV injection. Regenerating axons were visualized by injecting Alexa Fluor 594-conjugated cholera toxin subunit B (CTB-Alexa594) at 2 days prior to dissection. We found that significant enhanced regeneration from AAV-5’UTRm- injected optic nerves comparing to control nerves (Fig. 7g). By calculating the estimated number of CTB-Alexa594-positive axons, we found that AAV-5’UTRm promoted axon regeneration in optic nerves as effectively as seen in the sciatic nerves (Fig. 7h). Notably, *Gpr151* mRNA was not upregulated from retina tissues after optic nerve crush injury by RT-qPCR analysis, although *Atf3*, a well-recognized injury-responsive gene and known to be induced both from the PNS and the CNS after injury was dramatically upregulated (Fig. 7i). This result suggested that the upstream regulators responsible for *Gpr151* mRNA transcription was differently regulated in retina ganglion cells in response to nerve injury, while the downstream effectors of *Gpr151* mRNA was shared in the both systems^27^. These results suggested that 5’UTRm RNA was a suitable candidate for the therapeutic applications and a potential tool to study the mechanisms of axon regeneration from different nervous systems.

## Discussion

Neuronal responses after injury have been investigated using various experimental tools to identify mechanisms regulating the neuron-intrinsic regeneration program^1, 8, 55, 56^. Omics technologies have facilitated the gaining of new knowledge in the axonal regeneration pathway^57^. For example, DEG analysis is often used as a method to identify genes regulating the neuronal injury responses, based on the premise that significantly upregulated or downregulated genes may be functionally important for the biological process ^9^. Indeed, studies of transcriptomic profiling in injured peripheral neurons have uncovered injury-responsive genes, among which a subset has been implicated in the regulation of axon regeneration. However, there still remain many injury-induced genes without clearly defined function in axonal regeneration or other injury responses. To overcome the limitation of single-omics approach and enhance selectivity for interesting features of genes, we take combinatorial methods. There are earlier studies taking combinatorial approach comparing injury-induced proteome and transcriptome to identify signaling networks that are required for activation of the transcriptional regenerative program^57–59^. In the present study, we compare transcriptome of bulk-seq and ribosome-association efficiency of protein-coding DEGs after sciatic nerve injury and identify a group of genes that are upregulated in response to axonal injury without changes in their ribosome-association efficiency, supporting a striking discordance between transcriptional and translational controls after injury. The method in the current work of the comparative analysis between bulk-seq and *RiboTag*-seq has some limitations. First, if some bulk-seq mRNA largely origin from non-neuronal cells that do not express *Adv-Cre*, their upregulation shown in the bulk-seq may mainly represent injury responses in the non-neuronal cells. Second, both methods were performed with DRG tissues with only minimal axons, so local translation in axons was unexplored in this comparative study. This exciting uncovered area can be studied in the future using single-cell sequencing technique and axonal *RiboTag* experiment^60–65^.

*Gpr151* is one of the injury-responsive genes whose dramatic upregulation has consistently been reported in several DEG studies after injury from different nervous tissues^11, 26,27,28, 32, 66–69, 70^. GPR151 protein is known to play a molecular role as an orphan GPCR in signaling pathways controlling addiction and pain^25–27, 30, 31, 71–75^. Therefore, an easily driven hypothesis would be that the GPCR function of *Gpr151* is enhanced by the increased gene expression and contributes to the regulation of the regenerative signaling after injury. However, in contrast to this notion, we find that the upregulated *Gpr151* mRNA is not directed to ribosome complexes from our *RiboTag*-sequencing analysis. Instead of being recruited to ribosomes, upregulated *Gpr151* mRNA is associated with RBPs such as CSDE1. *Gpr151* mRNA binds to CSDE1 through a consensus binding sequence conserved in rodent and human genes in its 5’UTR and modulates the pool of CSDE1-binding mRNA.

GPR151 protein levels display no increase in mouse DRG tissue when axonal injury was introduced. Instead, the protein levels show a significant reduction after injury, indicating that additional mechanisms for regulating the GPR151 protein levels should be considered. In addition, GPR151 protein may have a negative role in axonal growth because overexpressing GPR151 protein inhibits neurite outgrowth in vitro. The enhanced axon regeneration observed in the sciatic nerves of GPR151 protein-null mutant mice also supports the inhibitory function of the GPR151 protein. A recent study by Antolin-Fontes and colleagues has demonstrated that GPR151 is constitutively active and interacts with the G-alpha inhibitory subunit Gαo1 in habenular neurons and that their interaction inhibits adenylyl cyclase, leading to a reduction in the cyclic AMP (cAMP) levels^25^. Since cAMP levels and the downstream PKA pathway are major components in the intrinsic axon regeneration program, GPR151 protein-dependent signaling may play the putative inhibitory role by suppressing the cAMP-PKA pathway in regenerating peripheral neurons. Because blocking cAMP-PKA pathway leads to activation of RhoA GTPase that is a crucial factor responsible for growth cone collapse, GPR151 protein overexpression may reduce the rates of axonal growth^76–79^. Furthermore, we speculate that the reduced GPR151 protein levels observed after nerve injury may be an adaptive response in the DRG neurons to circumvent the inhibitory effect of the protein function-dependent GPR151 pathway. This may be a possible mechanism for inefficient neurite projection from GPR151 protein-overexpressing DRG neurons in vitro. In addition, the non-existence of GPR151 protein may be responsible for the enhanced axon regeneration from GPR151 protein-null (KO) mice. Therefore, the role of GPR151 protein in axon regeneration and other injury responses is an interesting topic in future studies.

RBPs are implicated in diverse events in the nervous system such as neurodevelopment and memory consolidation and also involved in the pathogenesis of neurodegenerative diseases^80–83^. While the main role of RBPs is to regulate the fate of interacting RNA, the functions of RBPs are conversely modulated by interacting with specific regulatory RNAs ^18^. Hence, the molecular networks of transcripts and their interacting RBPs need to be delineated in order to correctly understand functional outcome of transcriptomic changes in a specific cellular response. Although we focus on CSDE1 as an interacting RBP of *Gpr151*-5’UTR, there are other RBPs identified in the binding protein-screening that putatively interact with the 5’UTR. For example, the interaction of the 5’UTR with NonO is confirmed by pull-down assays where the binding affinity is unaffected by the competitor containing the CSDE1-specific binding sequence in *Gpr151*-5’UTR. Therefore, it appears that NonO recognizes different sequences or structures of the 5’UTR from that interacting with CSDE1. As NonO is a nuclear RNA-binding protein and known to nuclear RNA via its double stranded structures for regulating RNA retention^52, 84^, *Gpr151* mRNA may interact with NonO in the nucleus before the mRNA leaves out the nucleus out through the double-stranded stem and loop structure that is not the CSDE1-interacting region. Importantly, overexpressing 5’UTR of *Gpr151* fails to further promote neurite outgrowth when CSDE1 is knocked down. Moreover, 5’UTR overexpression-induced growth potential is counterbalanced by CSDE1 overexpression and deletion of the CSDE1-binding motif abolishes the ability of 5’UTRm promoting neurite outgrowth. Therefore, we suggest that CSDE1 is a major functional target of *Gpr151*-5’UTR in axon regeneration, although it is unclear whether the interaction with NonO and other RBPs has regulatory roles in our current study.

Evidence builds for mRNA functioning as coding and non-coding RNAs (cncRNAs) with both protein-dependent and independent roles, although the role in axon-injury signaling has not been reported^22, 23^. In the present study, we identify *Gpr151* as a new cncRNAs in the axon regeneration pathway, based on the observation that its mRNA expression and ribosome-association is greatly uncoupled. As there are many other injury-responsive DEGs with the disconnected ribosomal targeting, more protein-coding genes may be functioning as cncRNAs involved in the regenerative response. Likewise, our understandings of the DEG functions in other cellular responses may be broadened by examining the potential non-coding roles of protein-coding mRNA as regulators of the interacting RBPs.

Our data show that expressing only the 5’UTR of *Gpr151* mRNA promotes peripheral axon regeneration. In addition, overexpressing the engineered form of 5’UTR for enhanced CSDE1-binding dramatically improves CNS axon regeneration in an optic nerve injury model. Therefore, we present the engineered 5’UTR of *Gpr151* as a potential tool for the manipulation of the regenerative potential of neurons. *Gpr151* mRNA is thus considered to be an RNA modulator that regulates CSDE1-RNA interaction through its 5’UTR and that has a function that is distinct from its protein in an axon regeneration paradigm.

## Acknowledgements

We thank Dr. Aaron DiAntonio (Washington University in St. Louis) and Dr. Valeria Cavalli (Washington University in St. Louis) for critical reading of the manuscript. This work was supported by the Samsung Research Funding & Incubation Center of Samsung Electronics under Project Number SRFC-MA1802-07.

## Author Contributions

Conceptualization, J.E.S., and Y.C.; Methodology, Formal Analysis, and Investigation, B.L., J.L., Y.J., E.S., Y.S., H.K., M.K., J.E.S., and Y.C.; Resources, Funding Acquisition, Y.C.; Supervision and Project Administration, J.E.S. and Y.C.; Data Curation and Visualization, J.E.S., and Y.C.; Writing – Original Draft, B.L., J.L., Y.J., J.E.S., and Y.C.; Writing – Review & Editing, J.E.S. and Y.C.

## Resource Availability

Further information and requests for reagents and resources should be directed to and will be fulfilled by the lead contact, Yongcheol Cho.

## Data and Code Availability

The data is available at NCBI Gene Expression Omnibus (GEO), accession GSE179145.

## Methods

### Antibodies

Anti-HA antibody (Abcam, ab9110; RRID: AB_307019), anti-GPR151 antibody (Aviva Systems Biology, OAAF06441), anti-α tubulin antibody (Santa Cruz Biotechnology, sc-53030; RRID: AB_2272440), anti-c-Jun antibody (Cell Signaling Technology, 9165; RRID: AB_2130165), anti-SCG10 antibody (Novus Biologicals, NBP1-49461; RRID: AB_10011569), anti-βIII tubulin antibody (TUJ1, Abcam, ab41489, RRID: AB_727049), anti-GAPDH antibody (Santa Cruz Biotechnology, sc-32233; RRID: AB_627679), anti-FLAG antibody (Cell Signaling Technology, 14793; RRID: AB_2572291).

### Animal models: (model organism: name used in paper: allele symbol)

Mouse: *RiboTag*: *Rpl22^tm^*^1^*^.1Psam^*; Mouse: *Adv*-Cre: *Avil^tm2(cre)Fawa^*; Mouse: CD-1: Crl:CD1(ICR); Mouse: C57BL/6J: C57BL/6J; Mouse: KO: *Gpr151^tm1Dgen^*. The *RiboTag* mouse carries a ribosomal protein gene with a floxed C-terminal exon followed by an identical exon tagged with hemagglutinin (HA) epitope. When *RiboTag* is crossed to a mouse expressing a cell-type-specific *Cre* recombinase, expression of the epitope-tagged protein is activated in the cell type of interest. A homozygote *RiboTag* mice (*Rpl22HA*) obtained from Jackson Laboratory and *Advillin*-Cre driver line ^85^ were crossed. Cre positive *RiboTag* animals (*Rpl22HA*+/+; *Adv*-Cre+) and the littermate control (*Rpl22HA*+/+; *Adv*-Cre-) mice were generated by crossing *Rpl22HA*+/+; *Adv*-Cre+/- male with *Rpl22HA*+/+ female. The KO mouse is the *Gpr151*-targeted mutant mouse (Deltagen) and carries a bacterial *LacZ* gene that is inserted into *Gpr151* gene and disrupt CDS. For in vivo animal studies, all the experiments were carried out in accordance with the Korea University Institutional Animal Care & Use Committee (KU-IACUC).

### Mouse surgery and DRG neurons cultures

Mice were anesthetized using 3% isoflurane in oxygen until unresponsive to toe pinch to confirm that the mouse is unconscious and kept on a heating pad for the duration of the surgery. The surgery site was shaved and disinfected using organic iodine of chlorhexidine solution. All surgery instruments were disinfected using autoclave and bead sterilizer. Mice were fed a diet (PicoLab 5053, Purina) and maintained on a 12 hours light/dark cycle (6 am to 6 pm). Male and female mice were housed in groups of up to 5 mice per cage. Age- and sex-matched male mice were used at the indicated age. Three-months or older mice were used for sciatic nerve surgery. Mice sciatic nerve was unilaterally exposed through a small incision made to the skin and muscles at mid-thigh level. Then, the sciatic nerve was crushed by forceps and the incision was closed by nylon suture. The animals were then subjected to post-operation care until euthanized for analysis.

To monitor neurite outgrowth of DRG neurons of adult mice, adult DRG cultures were prepared as described by Abe et al^86^. L4,5 DRGs from mice were collected with or without pre- conditioning by crushing the sciatic nerves before dissection and incubated in DMEM (Thermo Fisher Scientific, 41966029) with 0.7mg/ml Liberase TM (Roche, 5401119001), 0.6mg/ml DNase I (Sigma, 11284932001), and 10mg/ml bovine serum albumin (Sigma, A9418) for 15 min at 37’C. The DRG tissues were transferred to new tubes and incubated with trypsin-EDTA solution (Thermo Fisher Scientific, 12605010) for 15min at 37’C followed by trituration. Dissociated cells were plated on culture dishes coated with poly-d-lysine (Sigma, P0899) plus laminin (Thermo Fisher Scientific, 23017015) in DMEM with 10% fetal bovine serum (Thermo Fisher Scientific, 26140079), 1% penicillin-streptomycin, (Thermo Fisher Scientific, 15070063) and 1% Glutamax (Thermo Fisher Scientific, 35050061) and 50ng/ml nerve growth factor (Envigo, BT-5017) and incubated in humidified incubator at 37’C / 5% CO_2_.

### Purification of ribosome-associated RNA

Experimental procedures were adopted from Axon-Translating Ribosome Affinity Purification (TRAP) method with small changes ^87^. *RiboTag* x *Adv*-Cre mice (*RiboTag*; *Adv*-Cre+) were used for ribosome-associated RNA purification. *RiboTag*; *Adv*-Cre- was the counter mice as a negative control for immunoprecipitation. *RiboTag*; *Adv*-Cre+ mice expressed HA-epitope tagged RPL22 selectively from peripheral sensory neurons. L4,5 DRG tissues dissected from one *RiboTag*; *Adv*-Cre- female mouse (total four DRG tissues) were used for the negative control of the immunopurification. DRG tissues dissected from three *RiboTag*; *Adv*-Cre+ female mouse (all littermates, total six DRG tissues) were prepared from the left side of the mice without the left-side sciatic nerve crush injury. The L4,5 DRG right side of the same mice (total six DRG tissues) were dissected at 72 hours after the right-side sciatic nerve crush injury. Dissected DRG tissues were homogenized in lysis buffer; 1% NP-40, 20mM HEPES-KOH, 5mM MgCl_2_, 150mM KCl, 1mM DTT, SUPERase In (Thermo Fisher Scientific, AM2694), and Complete EDTA-free Protease Inhibitor Cocktail (Sigma, 11873580001) in the presence of cycloheximide (Sigma, C7698-1G), rapamycin (Sigma, R0395), and post-mitochondrial fractions were collected. The supernatant was pre-cleared by incubating 40μl of Dynabeads Protein A magnetic beads (Thermo Fisher Scientific, 10003D) at 4’C for 1 hour then transferred to a new collection tube. To immunoprecipitated HA-RPL22-associated ribosome complex, pre-cleared ribosome-mRNA complexes were immunoprecipitated by anti-HA antibody (Abcam, 9110) pre-conjugated with Dynabeads Protein A magnetic beads for 16 hours at 4’C with mild rotation. After 5 times washing, the immunoprecipitants were subjected to total RNA extraction using RNeasy micro kit (Qiagen, 74004) and in-column DNase I treatment for eliminating genomic DNA contaminants.

### Library preparation, RNA sequencing and analysis

RNA samples were prepared using RNAqueous Total RNA isolation kit (Thermo Fisher Scientific, AM1931). To analyze ribosome-associated RNA, Nanopore PCR-cDNA barcoding sequencing kit (Oxford Nanopore Technologies, SQK-PCB109) was used for library preparation. The quantified RNA samples (50ng) were subjected to reverse transcription reaction followed by strand-switching reaction as the manufacturer’s guide. The VN primer was annealed to the RNA to target poly A tail, and first-strand cDNA was synthesized by Maxima H Minus Reverse Transcriptase (Thermo Fisher Scientific, EP0752). The RNA-cDNA hybrid was purified using Agencourt AMPure XP magnetic beads (Beckman Coulter, A63880). The library from the Cre- negative control, Cre+ DRG without injury and Cre+ DRG with injury was subjected to barcoding process using barcode primers (Oxford Nanopore Technologies, BP01-BP12). PCR was performed using 2x LongAmp Taq Master Mix (New England Biolabs, M0287S) to select full-length transcripts followed by a second purification step using Agencourt beads (as described above). The barcoded libraries from the triplicates of the uninjured DRG tissue (labeled as u1, u2, u3) and from the triplicates of the injured DRG tissues (labeled as x1, x2, x3) were multiplexed with concentration normalization and loaded onto Flow Cell (Oxford Nanopore Technologies, FLO-MIN106D) as the manufacturer’s guide. The flow cells were used with more than 800 active pores from the initial pore check. *MinION* and *MinKNOW* (Oxford Nanopore Technologies, ver. 20.10.3) was used for sequencing with the integrated basecaller guppy for real-time fast-basecalling. Sequencing process was setup with duration of 72 hours, Q-score of 7, and fast-basecalling of *Guppy* (ver. 4.2.2). After basecalling and de-multiplexing from *MinKNOW*, the sequencing reads were subjected to *NanoStat* and *NanoPlot* to check quality of reads with scores and lengths ^88^, and mapped using *Minimap2* (option, -ax splice:hq -uf) ^89^ with a reference *Mus musculus* (GRCm38.p6, GCA_000001635.8). *Samtools* were used for sorting and indexing the mapped reads ^90^ and *edgeR* was used for statistical analysis of differential levels between uninjured and injured DRG samples ^36, 37^. To screen the target genes for downstream analysis, total 12,967 identified protein-coding genes were selected and plotted as x-y plot, with the x-axis of bulk-seq result indicating differential expression levels after sciatic nerve injury and the y-axis of the relative levels of ribosome-association efficiency after injury. To visualize the statistical significance from the edgeR analysis result, the relative level of -log_10_ *P*-value was indicated as the relative size of the circle.

To analyze total DEGs of bulk-seq. from adult DRG tissues after sciatic nerve crush injury, three C57/BL6 mice were used as a set of a biological replicate. Right side sciatic nerves were crushed and L4,5 DRGs were dissected at 24 hours after injury. Left side L4,5 DRGs were subjected to uninjured DRG samples. Total RNA was extracted from DRG tissues using RNAqueous Total RNA isolation kit (Thermo Fisher Scientific, AM1931). Nanopore libraries were prepared from total RNA by using Nanopore Direct RNA sequencing Kit (Oxford Nanopore Technologies, SQK-RNA002) according to the manufacturer’s instructions. The RT adapter was annealed to the RNA for reverse transcription and reverse-transcribed RNA was then purified using Agencourt RNAClean XP magnetic beads (Beckman Coulter, A63880). The purified RNA was eluted by adding 21μl Elution Buffer and resuspending the beads, incubating at room temperature for 10 min, pelleting the beads again, and transferring the supernatant (pre-sequencing mix) to a new tube. Then the sequencing adapter was ligated to the library. Nanopore sequencing was performed as per manufacturer’s guidelines using R9.4 SpotON flow cells (FLO-MIN106, ONT) with *MinION* sequencer winth *MinKNOW* software by setting duration 72 hours. Basecalled reads was mapped to the *Mus musculus* reference (GRCm38.p6, GCA_000001635.8) using *MiniMap2* (option, -ax splice -uf -k14). Indexing and statistical analysis was done using *Samtools* and *edgeR*. Total three biological replicates were subjected to *edgeR* analysis for testing statistical significance of injury-responsive differential expression.

To analyze CSDE1-associated RNA, mouse embryonic DRG spot-culture was used. Spot- culture DRG neurons were infected with control or 5’UTRm lentivirus at DIV2 and then lysed in TRAP lysis buffer at DIV5. The quantified total extracts were incubated with anti-CSDE1 antibody and the immunoprecipitants were subjected to RNA extraction and library preparation same as described in *RiboTag*-seq. analysis. Two biological replicates were subjected to *edgeR* analysis to test statistical significance.

### In vitro gene delivery by lentiviral transduction

Lentiviral transduction and spot-culture method was applied for highly efficient gene delivery to embryonic DRG cultures ^39, 68, 91–96^. Mouse embryo at E13.5 was used to collect DRG tissues. The dissociated DRGs were trypsinized (Thermo Fisher Scientific, 25300054) and plated on culture dishes coated with poly-D-lysine (Sigma, P0899) and laminin (Thermo Fisher Scientific, 23017015) as previously described ^91, 97^. The culture medium was based on Neurobasal (Thermo Fisher Scientific, 21103049) supplemented with 2% B-27 (Thermo Fisher Scientific, 17504044) including 1% Glutamax (Thermo Fisher Scientific, 35050061), 1 μM 5- fluoro-2′-deoxyuridine (Sigma, F0503), 1 μM uridine (Sigma, U3003), 1% penicillin–streptomycin (Thermo Fisher Scientific, 15070063), and 50 ng/mL 2.5S nerve growth factor (Envigo, BT-5017).

To maximize the efficiency of virus transduction, the spot-culture method was applied ^96^. The prepared suspension of embryonic DRG neurons was concentrated with density of 40 DRGs in 50 μl culture medium and the cell numbers were counted. The high dense embryonic DRG neurons were then plated on completely dried culture dishes that were pre-coated with poly-d-lysine plus laminin ^92^. The plating volume was 2.5 to 4 μl so to make a single spot on culture dishes. The spotted culture dishes were incubated in 37’C / 5% CO_2_ incubator for 20 min and then culture medium was gently added to the dishes. The medium added dishes were incubated in the incubator and lentivirus was transduced at DIV2. The spot-cultures were avoided from any agitations or movements.

To knock down or overexpress the indicated targets, lentivirus transduction was used with the spot-culture method that give the high efficiency of in vitro gene delivery from embryonic DRG neurons. pLKO (human U6 promoter) and FUGW (human Ubiquitin-C promoter) lentiviral constructing vector were used for polymerase III-dependent and polymerase II-dependent gene expression, respectively. To produce high titer lentiviral particles, Lenti-X Packaging Single Shot (Qiagen, 631276) was used for viral packaging then concentrated using Lenti-X Concentrator (Qiagen, 631232). The titer of the individual lentiviral solution was quantified by Lenti-X p24 Rapid Titer Kit (Qiagen, 632200) at the first round of production and then Lenti-X GoStix Plus Kit (Qiagen, 631280) was used for routine checks after the initial quantification.

To knock down mouse *Gpr151* from embryonic DRG neurons, the target sequence of TRCN0000026286 from TRC library database of BROAD Institute was sub-cloned into pLKO vector using AgeI/EcoRI restriction sites. TRCN0000026286 is 3’UTR-targeting shRNA for mouse *Gpr151* with target sequence of “GCAAAGATTTCTGCTTTCAAA”. To knockdown mouse *Csde1*, the target sequence of TRCN0000276864 that is also targeting the 3’UTR of mouse *Csde1* was sub-cloned into pLKO vector using AgeI/EcoRI restriction sites and the target sequence is “GCGCCTTATAGTGTCAATTTA”. The knockdown efficiency at RNA levels were determined by RT-qPCR analysis. To overexpress GPR151 proteins from the different forms of mRNA, three types of the GPR151 CDS-related inserts were sub-cloned into FUGW vector using In-Fusion Cloning kit (Takara, 638910). For control experimental sets, non-human and mouse targeting scrambled sequence “CCTAAGGTTAAGTCGCCCTCG” from *VectorBuilder* (VB180117-1020znr) was used.

To overexpress GPR151 protein from the mRNA that has CDS but no UTRs (FUGW- CDS), mouse *Gpr151* ORF clone (Origene, MR223707, NM_181543, Myc-DDK-tagged) was sub-cloned into FUGW vector including its original Myc-DDK-tag with no any UTRs at AgeI/EcoRI restriction sites. To overexpress GPR151 protein from the mRNA that has CDS and 5’UTR (FUGW-5’UTR-CDSwt), the 5’UTR sequence of mouse *Gpr151* (NM_181543) “CCAACCTAAACAAGAAGCTACCATCTGCAGGGAGGAGCTTG” was commercially synthesized from Integrated DNA Technologies (IDT) and inserted into AgeI restriction site at the upstream to the CDS of *Gpr151* using In-Fusion Cloning kit. To overexpress *Gpr151* mRNA that has the intact 5’UTR but has the mutation at the start codon of AUG to AUC (FUGW-CDS-mut), FUGW-5’UTR-CDSwt was subjected to PCR-based site-directed mutagenesis to introduce the point mutation using mutagenesis primer sets, 5’-GCTTGACCGGTATCGGAAAG GCAATGC-3’ and 5’-GCATT GCCTTTCCGATACCGGTCAAGC-3’ followed by Saner sequencing to confirm the mutation.

To overexpress 5’UTR RNA_WT_ or 5’UTRm RNA from embryonic DRG neurons, the oligonucleotides of 5’UTR sequences, “CCAACCTAAACAAGAAGCTACCATCTGCAGGGAGGAGCTTG” or 5’UTRm sequence, “CCAACCTAAACAAGAAGCTACCATCTGCAGGGAGG<u>TCGAA</u>G” were commercially synthesized from IDT and ligated into pLKO vector with AgeI/EcoRI restriction sites. To overexpress CSDE1, mouse *Csde1* ORF clone (Origene, MR210719, NM_144901, Myc-DDK-tagged) was sub-cloned into FUGW vector including its original Myc-DDK-tag using AgeI/EcoRI restriction sites.

### In vivo gene delivery by adeno-associated viral (AAV) transduction

To express 5’UTR_WT_ or 5’UTRm in adult mouse DRG neurons in vivo, adeno-associated virus (AAV, serotype 9) was delivered to neonatal CD-1 mice by facial vein injection using a Hamilton syringe (Hamilton, 1710 syringe with 33G/0.75-inch small hub removable needle). 10μl of AAV vrius with titer 10^13^ GC/ml injected mice were subjected to sciatic nerve crush injury at 4 to 6 weeks. The neuronal expression of GFP in adult DRG tissues after sacrificing the mice was validated from dissociated and cultured DRG neurons to visualize GFP signal in vitro. To express 5’UTR_WT_ or 5’UTRm in adult mouse retina ganglion cells in vivo, 1.5 ul of AAV was injected with pulled glass needle to vitreous of eye. The conjunctiva from the orbital part of the eye was cleared in order to expose the optic nerve, which was crushed for 3 seconds with Dumont #5 forceps (Fine Science Tools, 11254-20) and special care was taken not to damage the vein sinus. To avoid desiccation of the eye, a saline solution was applied before and after optic nerve crush. For cholera toxin b (CTB) axon tracing, 1% atropine sulfate solution (Bauch & Lomb) was applied to the eye to induce pupil dilation. A pulled glass needle was introduced in the vitreous through the ora serrate and 2μl of 2μg/μl Alexa Fluor 594-conjugated cholera toxin B (CTB) (Thermo Fisher Scientific, C34777) was injected. For optic nerve fixation and sectioning, mice were sacrificed at two days after CTB injection and eyes-optic nerves altogether were fixed by immersion in 4% paraformaldehyde solution for 2 hours. After being washed three times in PBS, eyes were transferred to 30% sucrose solution for 24 hours at 4’C. Optic nerves were then dissected out with micro scissors (Fine Science Tools, 15070-08), sectioned at 11μm in the cryostat and mounted in mounting medium ProLong Gold (Thermo Fisher Scientific, P36931).

### Assessment of in vivo axon regeneration from sciatic nerves or optic nerves

For sciatic nerves regeneration experiment, the right sciatic nerve was crushed by forceps and L4,5 DRGs were then dissected from spinal cord and subjected to downstream analysis such as immunohistochemistry, western blot analysis, RNA extraction, adult DRG cultures and fluorescence in situ hybridization analysis. Sciatic nerves were dissected at 3 days after crush injury and dissected, sectioned and immunostained with anti-SCG10 antibody and βIII tubulin antibody. SCG10 is a marker protein selectively labeling regenerating axons in PNS and is immediately degraded from the distal part of the axotomized axons after introducing injury ^39, 98^. To make the full visualization of the sciatic nerve sections with SCG10-immunostained regenerating axons in a single panel image, multiple fields of 10X objective-acquired raw images were taken along the nerve section and montaged automatically in microscope (EVOS FL Auto Cell Imaging System, Thermo Fisher Scientific, AMC1000). SCG10 fluorescence intensity was measured along the length of the nerve sections by making measuring windows of 100-pixel width (equivalent to 87.7μm) at every 100-pixel from the injury site to the distal part using *ImageJ* software ^40^. The acquired SCG10 intensities at every 100-pixel from the sections were then normalized to the crush site that was determined by TUJ1-counterstaining that clearly shows morphological deformation of the tissues by crushing with forceps and gave the maximal SCG10 intensity correlating with βIII tubulin-immunostaining where deformation of the nerve and axonal disruptions were visualized. Then, the normalized SCG10 intensity at the injury site of the maximal SCG10 intensity was set to one and presented as a x-y line plot as well as subjected to calculate regeneration index ^99^. Regeneration index was defined by the distance from the crush site to the location where normalized SCG10 intensity specified from in x-axis as 50% and 20%^100^.

For optic nerves regeneration experiment, the optic nerve sections were imaged with EVOS FL Auto Cell Imaging System in the same way for sciatic nerve imaging. From the acquired full panel of nerve sections, numbers of regenerating axons at different distances from the injury site were estimated following the published protocol described ^101–103^. The numbers of regenerating axons crossing lines set at 0.2, 0.5, 0.8 and 1mm from the injury site were quantified in 5 different slices per optic nerve using ImageJ. The cross-sectional width of the optic nerve was used to estimate the number of regenerating axons per μm. The total number of regenerating axons per each optic nerve at certain distance from the injury site (Σa_d_) was estimated using the following formula: Σa_d_ = π x (optic nerve radius)^2^ x [average axons/μm] / 11μm. For statistical analysis, the average numbers of regenerating axons from control and 5’UTRm-expressing mice at each distance were compared and the significance was analyzed by one-way ANOVA with Bonferroni post-test ^102^.

### Fluorescent in situ hybridization and immunohistochemistry

*Gpr151* mRNA was detected by a pair of target probe (RNAscope Probe-Mm-*Gpr151*, ACD, 317321) following the manufacturer’s instructions. L4,5 DRG tissues were fixed in 4% paraformaldehyde and permeabilized. The fixed sample tissues were cryo-blocked with OCT compound and sectioned at a 10μm thickness with cryotome. The sections were dehydrated with 50%, 70% to 100% Ethanol for 5min each at room temperature, followed by air dry for 5 min at room temperature. Tissue section was incubated in Protease 4 (ACD, from the kit RNAscope, 322000) solution for 15 min at room temperature. To wash, phosphate-buffered saline was applied five times to the tissue section. Target probe was pre-warmed at 40’C and then the probe was applied to tissue section and incubated for 2 hours in 40’C. Slide was then washed by wash buffer (ACD, from the kit RNAscope, 322000) twice. The fluorescence probes Amp-1-FL, Amp-2- FL and Amp3-FL were applied to tissue section for 30 min at 40’C. Finally, Amp-4-FL was applied and incubated for 15 min at 40’C. The prepared samples were mounted and analyzed under confocal microscope (Zeiss, LSM700). For immunohistochemistry, mouse tissues were dissected and fixed in 4% paraformaldehyde (Biosesang, P2031) for 1 hour at room temperature and incubated in PBS with 40% sucrose at 4’C for 16 hours. The cryoprotected tissues were embedded in OCT compound (Tissue-Tek) in plastic molds and sectioned in cryostat with 10 μm-thickness. The samples were blocked in blocking solution (5% normal goat serum in 0.5% Triton X-100 in PBS) for 1 hour and then incubated with primary antibodies diluted in blocking solution over night at 4’C. Samples were washed twice with 0.1% Triton X-100 in PBS, incubated with secondary antibodies for 1 hour at room temperature, followed by rinse three times with 0.1% Triton X-100 in PBS. VectaShield mounting medium (Vector Laboratories, H1000 or H1200) was used to mount the coverslip.

### Western blot and RT-qPCR analysis

For RT-qPCR analysis, RevertAid Reverse Transcriptase (Thermo Fisher Scientific, EP0441) and PowerUP SYBR green master mix (Thermo Fisher Scientific, A25918) was used. QuantaStudio 3 real-time PCR machine (Thermo Fisher Scientific) was used for qPCR analysis. The relative levels of target RNAs were quantified by RT-qPCR analysis. The following primer sets were used for qPCR analysis (F, forward; R, reverse).

*Gpr151*, F: 5’-CTGGGTTTGCCGACACCAAT-3’; R: 5’-AGAGAGACGGAATGATGGTCC-3’ *Jun*, F: 5’-CCTTCTACGACGATGCCCTC-3’; R: 5’-GGTTCAAGGTCATGCTCTGTTT-3’ *Atf3*, F: 5’-CAGTCACCAAGTCTGAGGCG-3’; R: 5’-GTGTCCGTCCATTCTGAGC-3’ *Fabp7*, F: 5’-GGACACAATGCACATTCAAGAAC-3’; R: 5’-CCGAACCACAGACTTACAGTTT-3’ *Vim*, F: 5’-CGGCTGCGAGAGAAATTGC-3’; R: 5’-CCACTTTCCGTTCAAGGTCAAG-3 To measure relative expression levels of viral transduced 5’UTR_WT_ or 5’UTR_m_, total RNA was extracted at DIV5 from cultured embryonic DRG neurons using miRNeasy Micro Kit (QIAGEN, 217084) and subjected to polyadenylating reaction using *Escherichia coli* poly(A) polymerase (E-PAP) Poly(A) Tailing Kit (Thermo Fisher Scientific, AM1350) in buffer containing 0.1mM ATP, 0.25mM MnCl_2_ at 37’C for 1 hour, followed by cDNA synthesis. For reverse transcription, poly(A) tag primer, CAGGTGGTCCAGTTTTTTTTTTTTTTT was used ^104^. For qPCR analysis, forward primer “5’-CCAACCTAAACAAGAAGCTACCATCTG-3’” and reverse primer “5’- GGTCCAGTTTTTTTTTTTTTTTCAAGCTCCTC-3’” for 5’UTR_WT_ and “5’- GGTCCAGTTTTTTTTTTTTTTTCTTCGACCTC-3’” for 5’UTRm was used.

For western blot analysis, mouse DRG tissues were dissected with or without sciatic nerve crush injury. Protein lysates were prepared in Cell Lysis Buffer (Cell Signaling Technology, 9803) supplemented with Complete EDTA-free Protease Inhibitor Cocktail (Sigma, 11873580001). The supernatant of the protein lysates was recovered from centrifugation and quantified protein lysate samples were subjected to SDS-PAGE. The protein bands bound to antibodies were visualized using an ECL system or IR-dye using Leica Odyssey System.

### In vitro replating assay

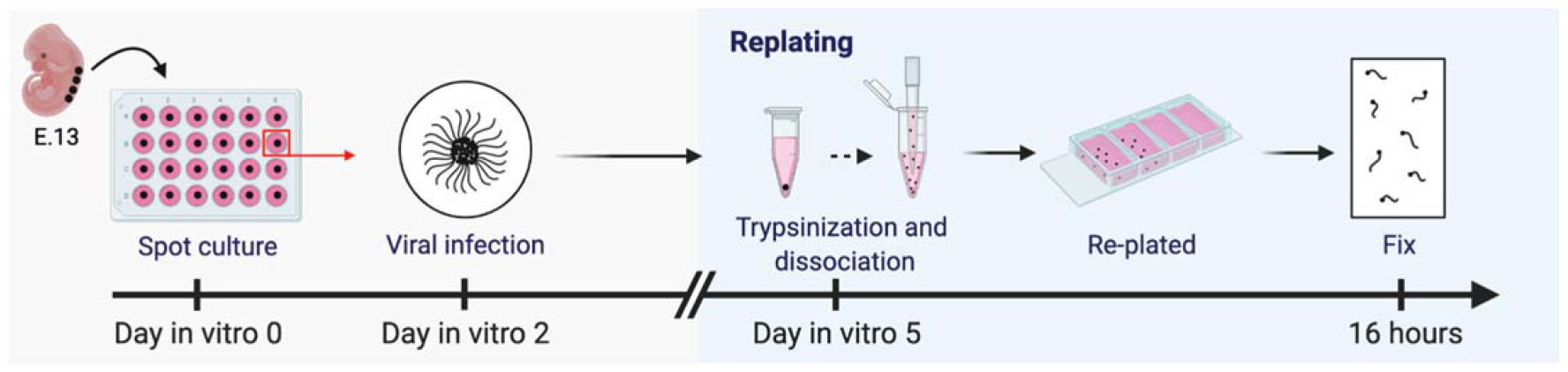

Primary embryonic DRG neuron cultures were prepared as spot-culture for viral transduction and replating assay. DRG tissues from mice at embryonic day 13.5 were dissociated in 0.05% trypsin-EDTA (Thermo Fisher Scientific, 25300054) and plated on poly-D-lysine (Sigma, P0899) / laminin (Thermo Fisher Scientific, 23017015)-coated dishes in Neurobasal medium (Thermo Fisher Scientific, 21103049) supplemented with 2% B-27 (Thermo Fisher Scientific, 17504044), 1% Glutamax (Thermo Fisher Scientific, 35050061), 1 μM 5-fluoro- 2’-deoxyuridine (Sigma, F0503), 1 μM uridine (Sigma, U3003), 1% penicillin-streptomycin (Thermo Fisher Scientific, 15070063), and 50 ng/ml 2.5S nerve growth factor (Envigo, BT-5017). The dissociated cell suspension was plated as spot-culture as described above. For in vitro gene delivery, spot-culture neurons were infected by lentivirus at DIV2. For replating assay, cultured neurons were replaced with DMEM (Hyclone, 500mlsh30243.01) / 0.05% Trypsin-EDTA mixture (1:1) at DIV 5. Neurons were incubated in 37°C, 5% CO_2_ incubator for 5 min. Then, neurons were pipetted with culture medium described above. Cells were then dissociated by gentle pipetting and transferred to new Lab-Tek chamber slides coated with poly-D-lysine/laminin. After re-plating, neurons were incubated for 12∼14 hours at 37°C with 5% CO2 then fixed in 4% paraformaldehyde (Biosesang, P2031) for 15min at room temperature. Neurite lengths were measured and assessed using the Neurite Tracer for ImageJ.

### Pull down assay and mass spectrometry analysis

All the RNA baits used for pull-down assay were commercially synthesized by IDT. Total protein lysates were prepared from DRG tissues dissected from 6-weeks old CD-1 mice, lysed in lysis buffer (1% NP-40, 20mM HEPES-KOH, 5mM MgCl_2_, 150mM KCl, 1mM DTT, SUPERase In (Thermo Fisher Scientific, AM2694), and Complete EDTA-free Protease Inhibitor Cocktail (Sigma, 11873580001)). 1 mg of protein lysates was subjected as an input for each pull-down condition. The quantified protein input was incubated with the indicated biotin-RNA baits for 16 hours at 4’C, followed by incubation with *Dynabeads MyOne* Streptavidin T1 (Thermo Fisher Scientific, 65601) for additional 1 hour at 4’C with rotation. The magnetic beads complexes were washed and recovered by magnet (Thermo Fisher Scientific, 12321D) as following the manufacturer’s instructions. Total protein from the precipitated magnetic beads were eluted by incubating at 95’C for 10 min in 1X SDS-PAGE sampling buffer then subjected to SDS-PAGE separation. For identifying the eluted proteins by mass spectrometry analysis, protein bands from PAGE gel were visualized by Coomassie staining method and sliced to subject to in-gel digestion. All the mass spectrometry analysis including sample preparation were done by YPRC (Yonsei Proteome Research Center, Seoul Korea).

For validating 5’UTR-CSDE1 interaction, biotin-conjugated RNA baits with 10 μM concentration were incubated in 200 μg total protein lysates prepared from cultured embryonic DRG neuron extracts in lysis buffer. After overnight incubation of biotin-RNA-lysates complexes at 4’C, *Dynabeads MyOne* Streptavidin T1 was applied and further incubated for 1 hour at 4’C. For competition pull-down assay, the competitor RNA with no biotin conjugation was co-incubated with bait RNA. The magnetic beads were washed using magnet and then subjected to SDS-PAGE analysis. For CSDE1-immunoprecipitation to prepare CSDE1-associated RNA after sciatic nerve injury in vivo, right side L4,5 DRG tissues from three C57/BL6 mice were dissected at 24 hours after sciatic nerve crush injury. The left side L4,5 DRG tissues of the same mice that had no pre-conditioning injury were used for total lysates of uninjured samples. Tissues were homogenized in TRAP lysis buffer as described above. CSDE1 was immunoprecipitated by anti- CSDE1 antibody (Abcam, ab201688) pre-bound to *Dynabeads* Protein G (Thermo Fisher Scientific, 10004D) from total 82 μg input lysates for 16 hours at 4’C. The precipitants were washed 5-times using *DynaMag-2* (Thermo Fisher Scientific, 12321D) and subjected to reverse transcription for quantitative real time PCR to quantify relative levels of CSDE1-associated *Gpr151*, *Vim*, and *Fabp7*.

## Supplementary information

**Supplementary Fig. 1.** Western blot analysis of L4,5 DRG tissues dissected from *RiboTag* x *Adv*-Cre mice with (+Injury) or without (-Injury) sciatic nerve crush. Anti-HA epitope antibody was used to detect RPL22HA protein.

**Supplementary Fig. 2.** Top 50 injury-responsive DEGs from edgeR analysis. *Gpr151* was highlighted and ranked 14th and 2nd in injury-responsive upregulation and expression level (LR of edgeR), respectively.

**Supplementary Fig. 3.** Adult L4,5 DRG neurons dissected from control or GPR151-protein null (KO) mice without sciatic nerve crush injury (-SNI), plated, cultured for 12 hours, fixed and immunostained with anti-βIII tubulin antibody. Scale bar, 100μm. The individual image represented a randomly selected raw microscopic field.

**Supplementary Fig. 4.** In vitro replating assay of embryonic DRG neurons. (a) The images were representative images of replated embryonic DRG neurons of control, *Gpr151* knockdown (KD), *Gpr151* knockdown + *Gpr151*-CDS overexpression (KD + CDS), *Gpr151* knockdown + 5’UTR of *Gpr151* overexpression (KD +5’UTR). (b) Average relative neurite length of (a). Statistical analysis from two biological replicates with cell numbers n=31+25, 30+30, 27+29, 32+23 for each condition; \**p*<0.05, ns, not significant by ANOVA followed by Tukey tests; mean±SEM.

**Supplementary Fig. 5.** The original microscopic field raw images for constructing the collage image of Fig. 4c, control.

**Supplementary Fig. 6.** The original microscopic field raw images for constructing the collage image of Fig. 4c, KD.

**Supplementary Fig. 7.** The original microscopic field raw images for constructing the collage image of Fig. 4c, KD+5’UTR-CDSwt.

**Supplementary Fig. 8.** The original microscopic field raw images for constructing the collage image of Fig. 4c, KD+5’UTR-CDSmut.

**Supplementary Fig. 9.** The original microscopic field raw images for constructing the collage image of Fig. 4h, control.

**Supplementary Fig. 10.** The original microscopic field raw images for constructing the collage image of Fig. 4h, KD.

**Supplementary Fig. 11.** The original microscopic field raw images for constructing the collage image of Fig. 4h, CDS.

**Supplementary Fig. 12.** The original microscopic field raw images for constructing the collage image of Fig. 4h, 5’UTR.

**Supplementary Fig. 13.** In vitro replating assay of embryonic DRG neurons. (a) Representative images of control, 5’UTRm-overexpressing, and 5’UTRmΔCSDE1-overexpressing embryonic DRG neurons. Scale bar, 100 μm. (b) Statistical analysis of regenerating neurite length of (a) (n=254, 198, 167 for control, shCSDE1 and shKHDRBS1; ***p<0.001, ns, not significant by ANOVA followed by Tukey tests; mean±SEM).

**Supplementary Fig. 14.** Lentivirus transduction efficiency from in vitro replating assay. Embryonic DRG neurons were prepared by spot-culture method and GFP-expressing control pLKO lentivirus was infected at DIV2. The neurons were replated at DIV5 and fixed and immunostained with βIII tubulin antibody. (a) Representative images of replated DRG neurons from three biological replicates. GFP fluorescence expressed under CMV-promoter of the 2nd ORF of lentivirus vector was visualized. Scale bar, 100 μm. (b) Statistical analysis of the cell numbers from (a). Box indicated 25% to 75% of the distribution. Closed box indicated mean.

**Supplementary Data 1.** Statistical analysis of injury-responsive DEGs from bulk-seq of adult mouse L4,5 and RiboTag-seq from edgeR analysis, for fig. 1.

**Supplementary Data 2.** The list of identified peptides by mass spectrophotometry analysis of fig. 5a.

**Supplementary Data 3.** Statistical analysis and gene ontology analysis of CSDE1- immunoprecipitation and sequencing result from control and 5’UTRm-overexpressing embryonic DRG neurons, for fig. 6.

