## Supplementary information 1 for "Promoting axon regeneration by enhancing the non-coding function of the injury-responsive coding gene *Gpr151*"

### Supplementary Information
for

**Promoting axon regeneration by enhancing the non-coding function of the injury-responsive coding gene *Gpr151***

**Bohm Lee[#1], Jinyoung Lee[#1], Yewon Jeon[#1], Eunsoo Jang[1], Yeonsoo Oh[1], Hyemin Kim[1], Minjae Kwon[1], Jung Eun Shin[1,2,3] and Yongcheol Cho*[1]**

[1] Department of Life Sciences, Korea University, Seoul 02841, Republic of Korea

[2] Department of Molecular Neuroscience Dong-A University College of Medicine, Busan 49201, Republic of Korea

[3] Department of Translational Biomedical Sciences, Graduate School of Dong-A University, Busan 49201, Republic of Korea

[#]These authors contributed equally to this work.

**This PDF file includes:**

Supplementary Figure 1 to 14

**Supplementary Fig. 1.**

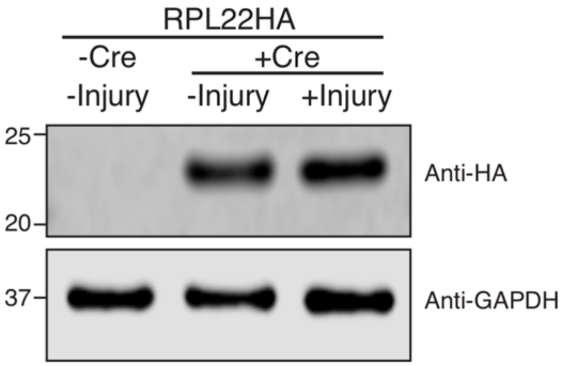


Western blot analysis of L4,5 DRG tissues dissected from *RiboTag* x *Adv*-Cre mice with (+Injury) or without (-Injury) sciatic nerve crush. Anti-HA epitope antibody was used to detect RPL22HA protein.

**Supplementary Fig. 2.**

| ENSG | Gene symbol | u1 | u2 | u3 | x1 | x2 | x3 | logFC | RANK | logCPM | RANK | LR | RANK | FDR |
|---|---|---|---|---|---|---|---|---|---|---|---|---|---|---|
| ENSMUSG00000024770 | Lipn | 0 | 0 | 0 | 12 | 10 | 11 | 10.9 | 1 | 2.5131 | 33 | 463 | 11 | 3.5394E-99 |
| ENSMUSG00000031635 | Anxa10 | 0 | 0 | 0 | 6 | 4 | 4 | 9.719 | 2 | 1.3699 | 36 | 240 | 25 | 3.75706E-51 |
| ENSMUSG00000038676 | Ucn | 0 | 0 | 0 | 4 | 5 | 5 | 9.628 | 3 | 1.283 | 41 | 257 | 23 | 1.16972E-54 |
| ENSMUSG00000050359 | Sprr1a | 1 | 0 | 0 | 434 | 275 | 343 | 9.537 | 4 | 7.4559 | 3 | 722 | 4 | 3.7442E-155 |
| ENSMUSG00000069355 | Gm5152 | 0 | 0 | 0 | 2 | 3 | 2 | 8.688 | 5 | 0.398 | 46 | 167 | 36 | 2.61306E-35 |
| ENSMUSG00000029819 | Npy | 1 | 0 | 0 | 120 | 76 | 85 | 7.543 | 6 | 5.5673 | 12 | 550 | 7 | 5.8063E-118 |
| ENSMUSG00000048764 | Tmprss11f | 0 | 0 | 0 | 4 | 3 | 4 | 7.337 | 7 | 0.9312 | 43 | 219 | 28 | 1.54387E-46 |
| ENSMUSG00000026247 | Ecel1 | 1 | 1 | 1 | 65 | 48 | 56 | 6.671 | 8 | 4.8399 | 19 | 717 | 5 | 3.8054E-154 |
| ENSMUSG00000030898 | Cckbr | 0 | 0 | 0 | 32 | 25 | 28 | 6.577 | 9 | 3.8505 | 24 | 650 | 6 | 1.1061E-139 |
| ENSMUSG00000023387 | Kcnk16 | 0 | 0 | 0 | 5 | 4 | 5 | 6.454 | 10 | 1.3398 | 38 | 254 | 24 | 3.6067E-54 |
| ENSMUSG00000028255 | Clca3 | 0 | 0 | 0 | 2 | 2 | 2 | 6.449 | 11 | 0.1113 | 49 | 125 | 43 | 2.38137E-26 |
| ENSMUSG00000049796 | Crh | 0 | 0 | 0 | 3 | 3 | 4 | 6.438 | 12 | 0.8411 | 44 | 191 | 30 | 1.63408E-40 |
| **ENSMUSG00000042816** | **Gpr151** | **1** | **1** | **1** | **90** | **87** | **95** | **6.168** | **13** | **5.5285** | **14** | **1091** | **2** | **7.2801E-235** |
| ENSMUSG00000034783 | Cd207 | 0 | 0 | 0 | 2 | 2 | 3 | 5.962 | 14 | 0.4024 | 45 | 136 | 39 | 1.21301E-28 |
| ENSMUSG00000028602 | Tnfrsf8 | 0 | 0 | 0 | 6 | 5 | 3 | 5.804 | 15 | 1.367 | 37 | 190 | 31 | 2.24906E-40 |
| ENSMUSG00000026628 | Atf3 | 8 | 7 | 9 | 428 | 363 | 417 | 5.679 | 16 | 7.682 | 2 | 1190 | 1 | 2.7383E-256 |
| ENSMUSG00000076434 | Wfdc3 | 0 | 0 | 0 | 9 | 11 | 13 | 5.267 | 17 | 2.5845 | 32 | 308 | 22 | 1.11915E-65 |
| ENSMUSG00000024907 | Gal | 2 | 2 | 3 | 82 | 72 | 84 | 5.113 | 18 | 5.3582 | 16 | 794 | 3 | 1.6564E-170 |
| ENSMUSG00000031074 | Fgf3 | 0 | 0 | 1 | 14 | 15 | 15 | 5.08 | 19 | 2.9159 | 30 | 454 | 12 | 3.9194E-97 |
| ENSMUSG00000020838 | Slc6a4 | 0 | 1 | 2 | 23 | 24 | 24 | 4.56 | 20 | 3.6362 | 26 | 158 | 37 | 1.70925E-33 |
| ENSMUSG00000019890 | Nts | 4 | 4 | 5 | 97 | 79 | 93 | 4.412 | 21 | 5.561 | 13 | 493 | 10 | 1.2475E-105 |
| ENSMUSG00000039364 | Sectm1b | 0 | 0 | 1 | 7 | 8 | 10 | 4.024 | 22 | 2.1725 | 34 | 215 | 29 | 9.19377E-46 |
| ENSMUSG00000073680 | Tmem88b | 1 | 1 | 1 | 19 | 15 | 18 | 3.862 | 23 | 3.2222 | 27 | 317 | 20 | 1.31799E-67 |
| ENSMUSG00000026639 | Lamb3 | 0 | 0 | 1 | 4 | 3 | 4 | 3.757 | 24 | 1.1088 | 42 | 133 | 40 | 4.85209E-28 |
| ENSMUSG00000035202 | Lars2 | 1475 | 1274 | 243 | 691 | 840 | 38740 | 3.751 | 25 | 12.816 | 1 | 10.9 | 50 | 0.011747928 |
| ENSMUSG00000051379 | Flrt3 | 20 | 18 | 20 | 289 | 217 | 254 | 3.716 | 26 | 7.0935 | 6 | 494 | 9 | 9.0682E-106 |
| ENSMUSG00000043091 | Tuba1c | 3 | 5 | 6 | 58 | 97 | 24 | 3.69 | 27 | 5.0106 | 17 | 68.6 | 48 | 2.04786E-14 |
| ENSMUSG00000011008 | Mcoln2 | 0 | 0 | 0 | 4 | 5 | 4 | 3.555 | 28 | 1.3348 | 39 | 153 | 38 | 2.00724E-32 |
| ENSMUSG00000027261 | Hao1 | 0 | 0 | 0 | 2 | 2 | 2 | 3.52 | 29 | 0.3232 | 47 | 93.4 | 45 | 1.20517E-19 |
| ENSMUSG00000058153 | Sez6l | 13 | 13 | 13 | 171 | 126 | 149 | 3.518 | 30 | 6.3392 | 8 | 443 | 14 | 7.89918E-95 |
| ENSMUSG00000063632 | Sox11 | 4 | 5 | 6 | 57 | 53 | 58 | 3.472 | 31 | 4.9373 | 18 | 412 | 15 | 3.75469E-88 |
| ENSMUSG00000035498 | Cdcp1 | 0 | 0 | 0 | 2 | 2 | 2 | 3.456 | 32 | 0.2702 | 48 | 83.3 | 46 | 1.65047E-17 |
| ENSMUSG00000026834 | Acvr1c | 1 | 1 | 2 | 12 | 12 | 13 | 3.427 | 33 | 2.7681 | 31 | 239 | 26 | 7.5412E-51 |
| ENSMUSG00000019647 | Sema6a | 30 | 28 | 25 | 344 | 237 | 281 | 3.377 | 34 | 7.3005 | 5 | 329 | 18 | 3.25707E-70 |
| ENSMUSG00000025473 | Adam8 | 7 | 7 | 9 | 84 | 73 | 76 | 3.348 | 35 | 5.4185 | 15 | 451 | 13 | 1.16088E-96 |
| ENSMUSG00000022044 | Stmn4 | 3 | 3 | 4 | 31 | 25 | 39 | 3.329 | 36 | 4.1355 | 22 | 219 | 27 | 1.47869E-46 |
| ENSMUSG00000001473 | Tubb6 | 13 | 13 | 14 | 117 | 110 | 121 | 3.115 | 37 | 6.0148 | 10 | 525 | 8 | 2.107E-112 |
| ENSMUSG00000046223 | Plaur | 3 | 3 | 5 | 31 | 25 | 32 | 3.084 | 38 | 4.0589 | 23 | 186 | 34 | 2.35722E-39 |
| ENSMUSG00000024883 | Rin1 | 2 | 1 | 2 | 16 | 10 | 14 | 3.074 | 39 | 2.9406 | 29 | 171 | 35 | 3.96221E-36 |
| ENSMUSG00000036390 | Gadd45a | 10 | 12 | 13 | 89 | 96 | 103 | 3.043 | 40 | 5.7554 | 11 | 388 | 16 | 4.7432E-83 |
| ENSMUSG00000044626 | Liph | 0 | 1 | 1 | 4 | 3 | 5 | 2.989 | 41 | 1.2947 | 40 | 116 | 44 | 1.52829E-24 |
| ENSMUSG00000037411 | Serpine1 | 3 | 2 | 4 | 28 | 21 | 20 | 2.98 | 42 | 3.6968 | 25 | 130 | 41 | 1.38014E-27 |
| ENSMUSG00000031574 | Star | 5 | 4 | 5 | 36 | 34 | 36 | 2.893 | 43 | 4.3362 | 21 | 363 | 17 | 1.40486E-77 |
| ENSMUSG00000009185 | Ccl8 | 1 | 0 | 1 | 5 | 9 | 6 | 2.875 | 44 | 1.8643 | 35 | 75.9 | 47 | 6.09955E-16 |
| ENSMUSG00000033060 | Lmo7 | 19 | 16 | 12 | 139 | 112 | 97 | 2.87 | 45 | 6.0481 | 9 | 186 | 33 | 1.88248E-39 |
| ENSMUSG00000014599 | Csf1 | 20 | 18 | 20 | 164 | 125 | 138 | 2.869 | 46 | 6.3422 | 7 | 322 | 19 | 1.00523E-68 |
| ENSMUSG00000078532 | Nkain1 | 2 | 2 | 3 | 13 | 19 | 14 | 2.851 | 47 | 3.1359 | 28 | 129 | 42 | 3.53562E-27 |
| ENSMUSG00000075249 | Fsip2 | 0 | 0 | 0 | 2 | 1 | 2 | 2.846 | 48 | 0.0732 | 50 | 59.7 | 49 | 1.50399E-12 |
| ENSMUSG00000044017 | Gpr133 | 7 | 6 | 7 | 55 | 46 | 44 | 2.827 | 49 | 4.7881 | 20 | 310 | 21 | 3.99524E-66 |
| ENSMUSG00000037428 | Vgf | 43 | 40 | 50 | 349 | 315 | 227 | 2.736 | 50 | 7.418 | 4 | 190 | 32 | 2.35293E-40 |

Top 50 injury-responsive DEGs from edgeR analysis. Gpr151 was highlighted and ranked 14 and 2 in injury-responsive upregulation and expression level (LR of edgeR), respectively.

**Supplementary Fig. 3.**

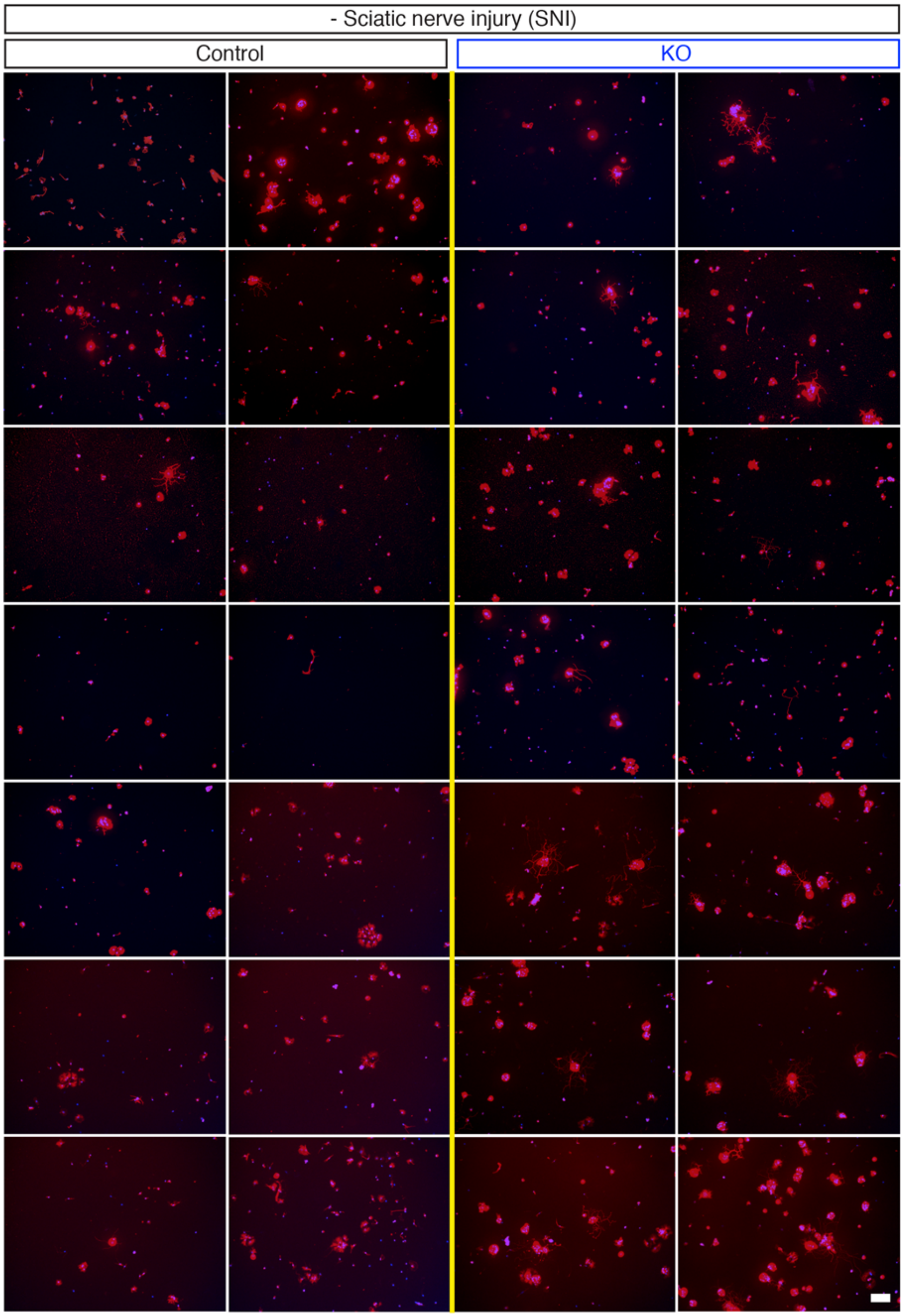


Adult L4,5 DRG neurons dissected from control or GPR151-protein null (KO) mice without sciatic nerve crush injury (-SNI), plated, cultured for 12 hours, fixed and immunostained with anti-βIII tubulin antibody. Scale bar, 100µm. The individual image represented a randomly selected raw microscopic field.

**Supplementary Fig. 4.**

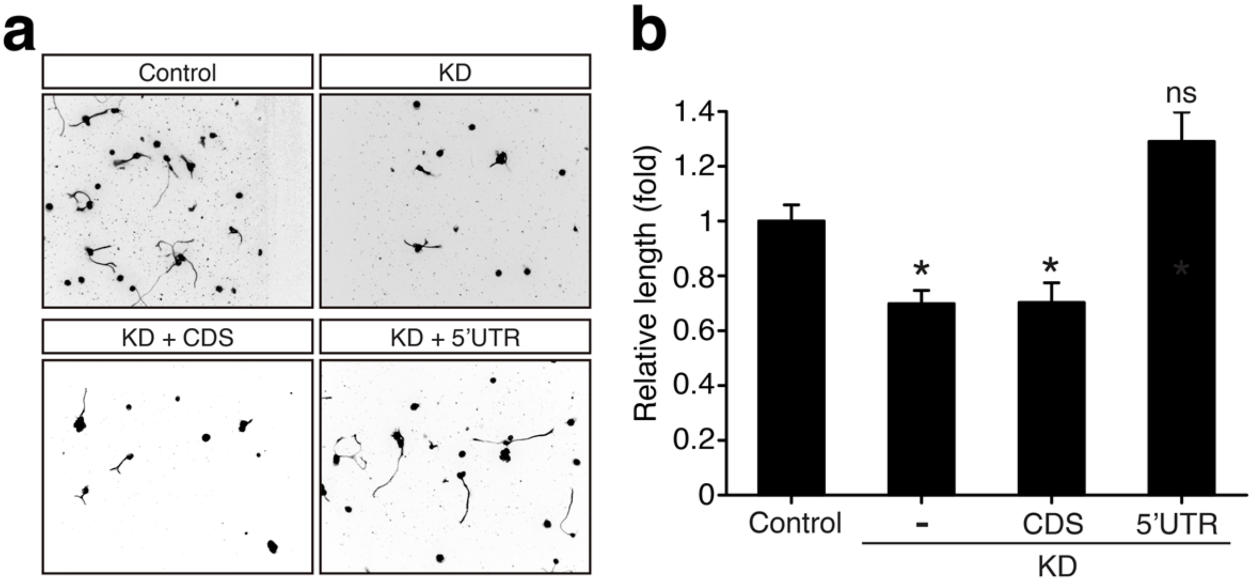


In vitro replating assay of embryonic DRG neurons. (a) The images were representative images of replated embryonic DRG neurons of control, *Gpr151* knockdown (KD), *Gpr151* knockdown + *Gpr151*-CDS overexpression (KD + CDS), *Gpr151* knockdown + 5’UTR of *Gpr151* overexpression (KD +5’UTR). (b) Average relative neurite length of (a). Statistical analysis from two biological replicates with cell numbers n=31+25, 30+30, 27+29, 32+23 for each condition; *$p$<0.05, ns, not significant by ANOVA followed by Tukey tests; mean±SEM.

**Supplementary Fig. 5.**

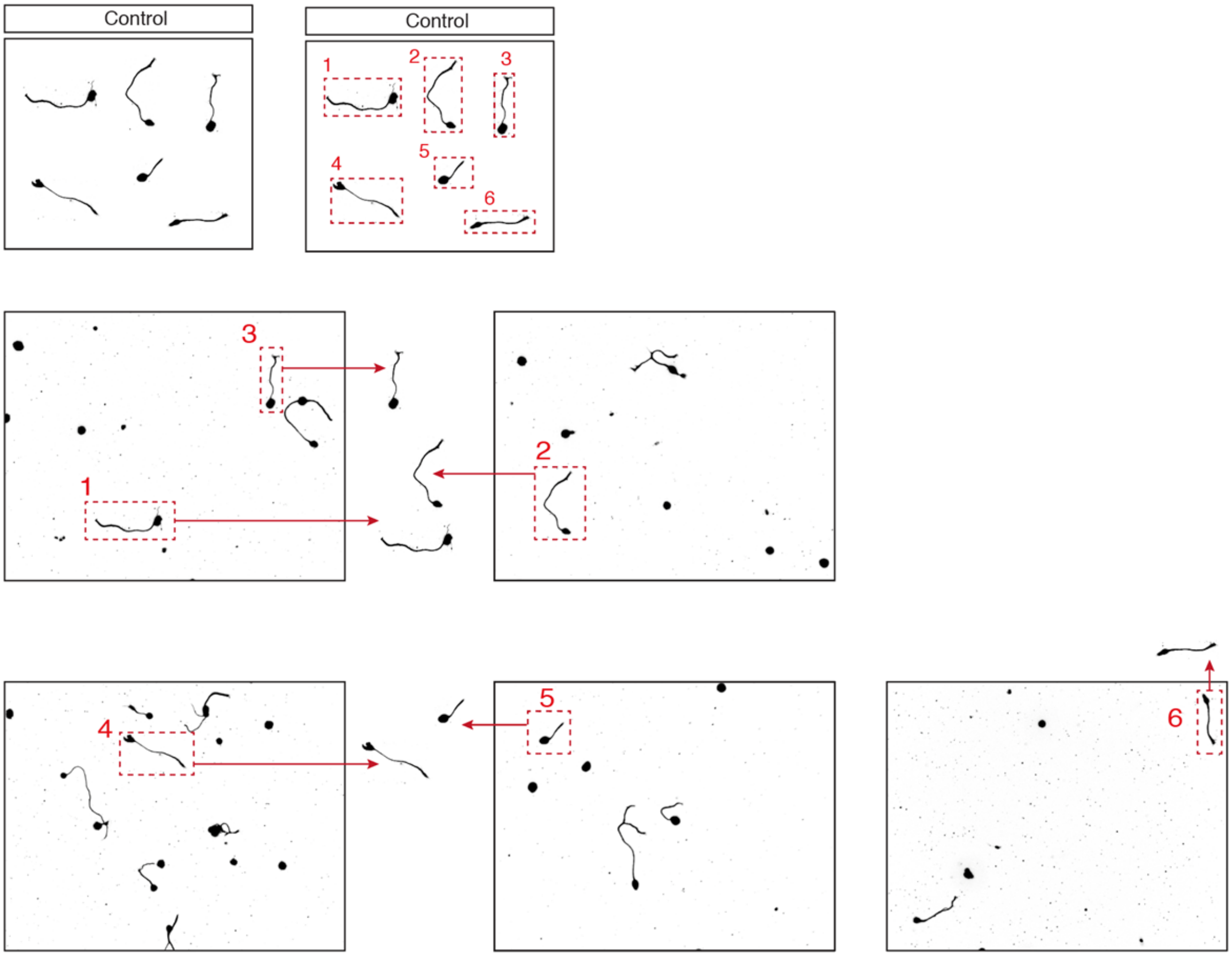


The original microscopic field raw images for constructing the collage image of Fig. 4c, control.

**Supplementary Fig. 6.**

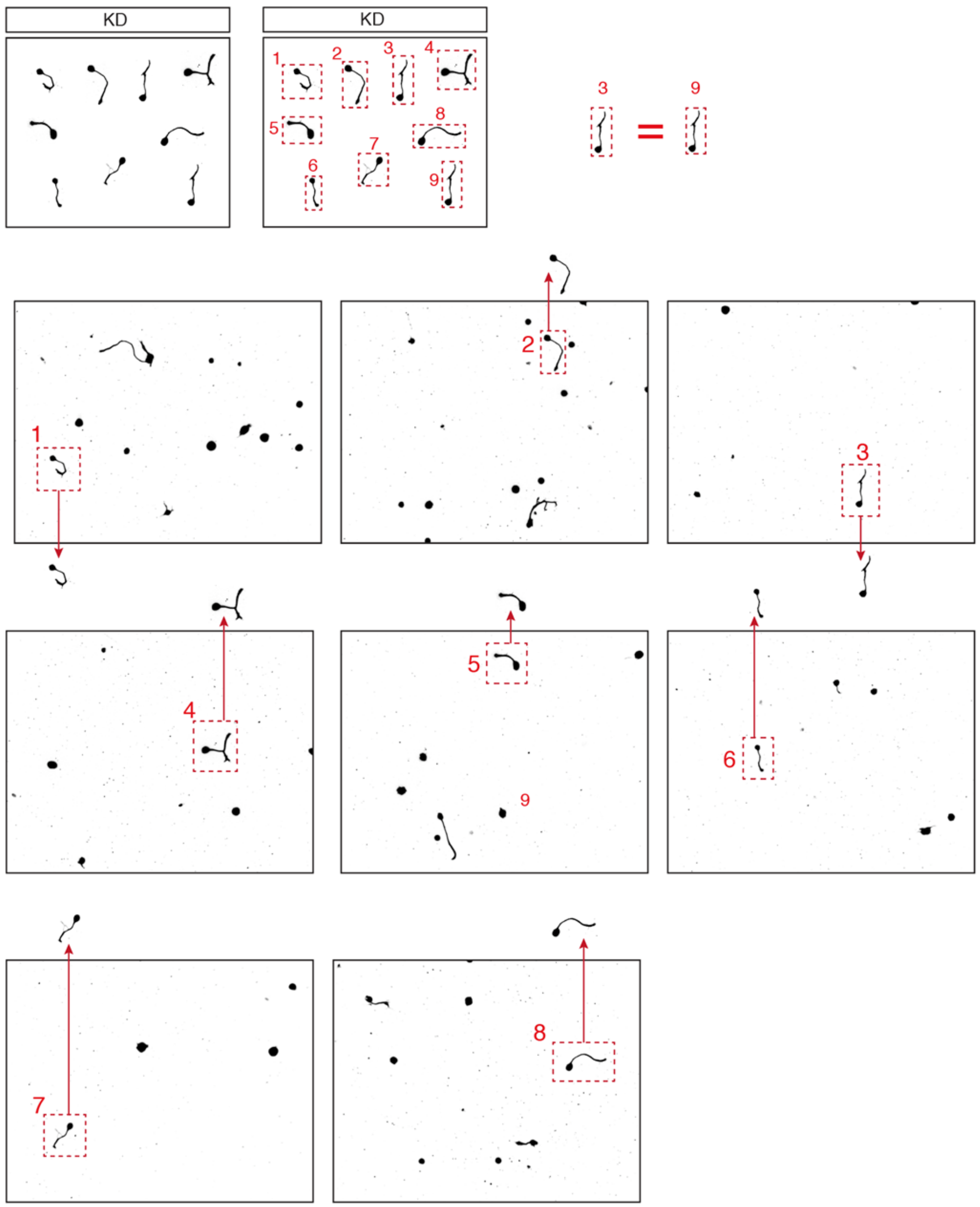


The original microscopic field raw images for constructing the collage image of Fig. 4c, KD.

**Supplementary Fig. 7.**

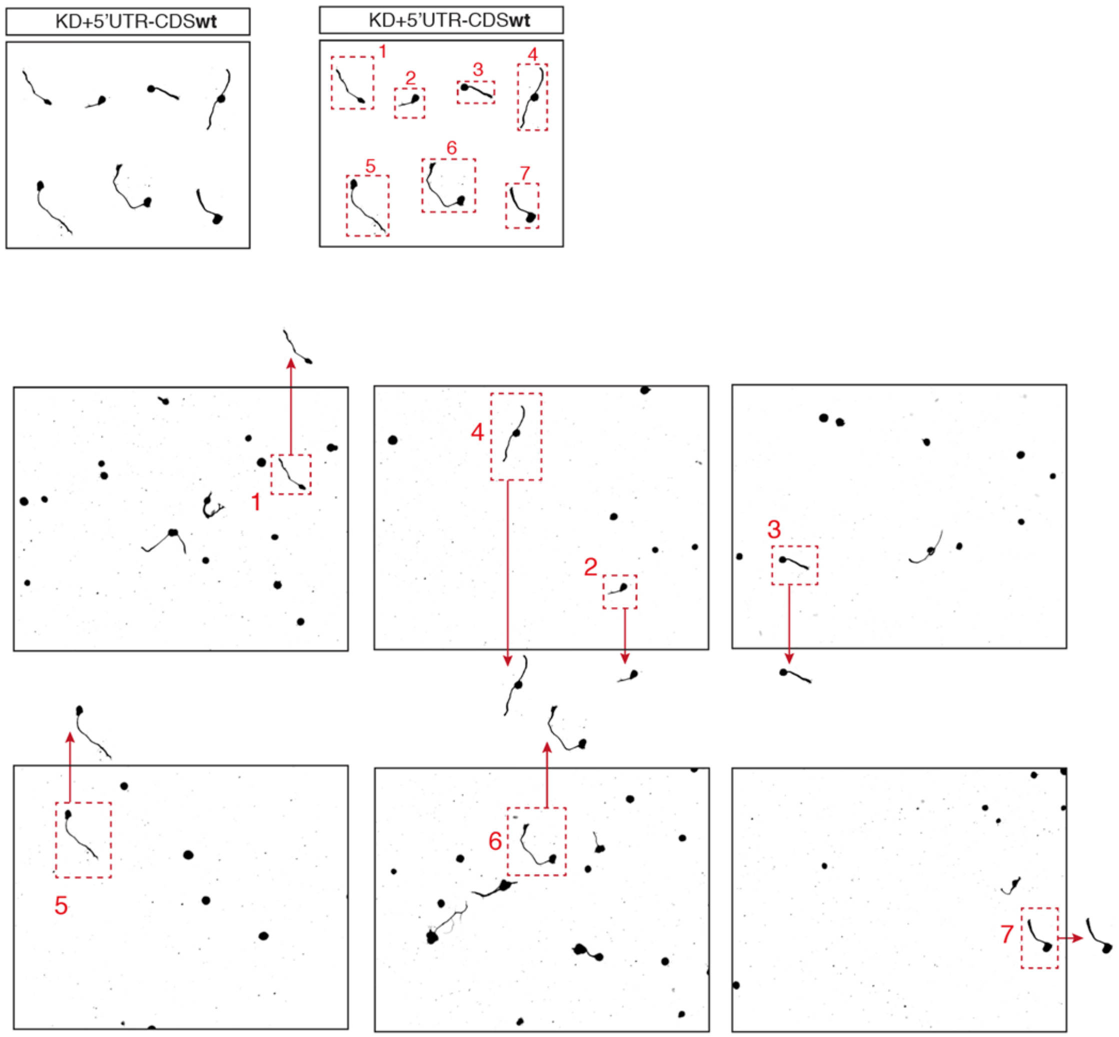


The original microscopic field raw images for constructing the collage image of Fig. 4c, KD+5'UTR-CDSwt.

**Supplementary Fig. 8.**

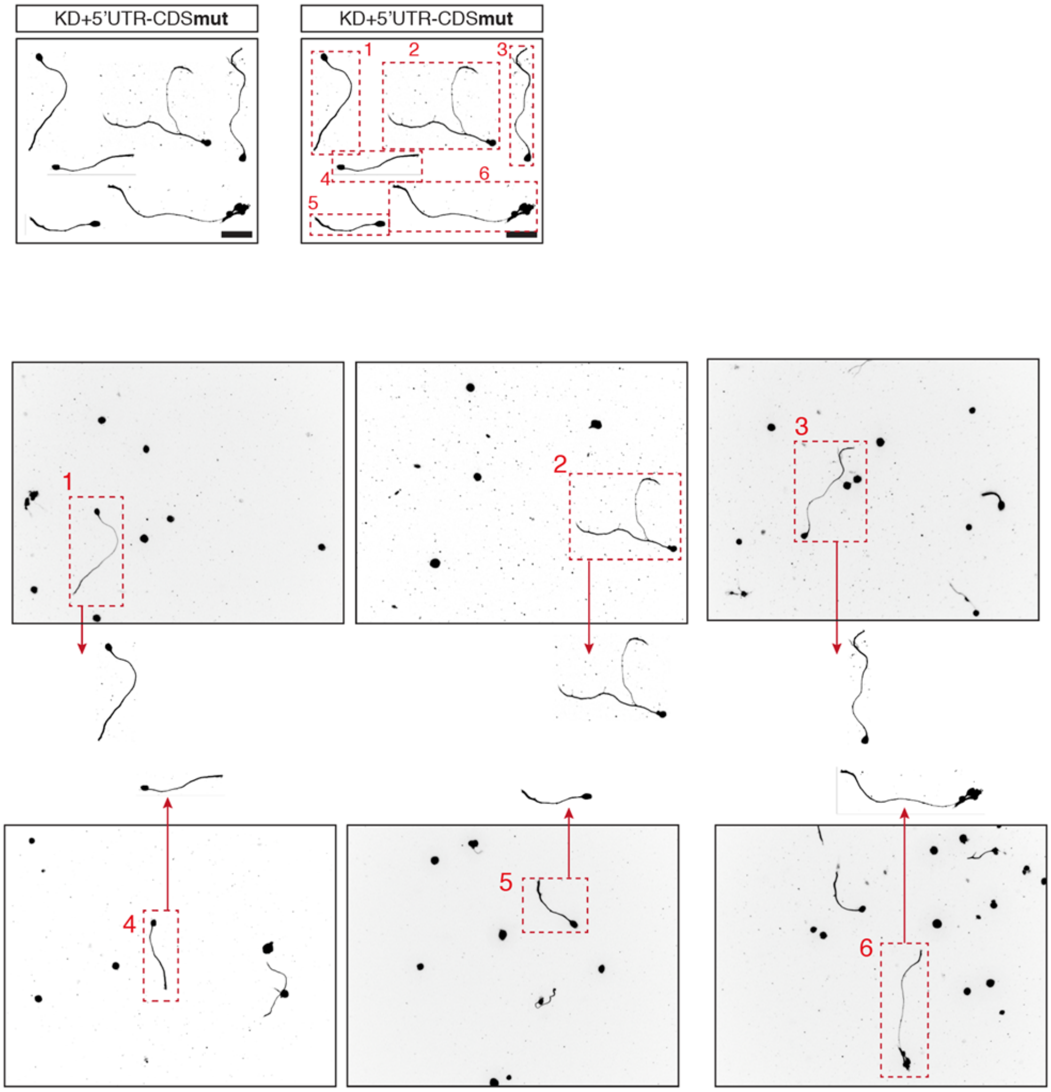


The original microscopic field raw images for constructing the collage image of Fig. 4c, KD+5’UTR-CDSmut.

**Supplementary Fig. 9.**

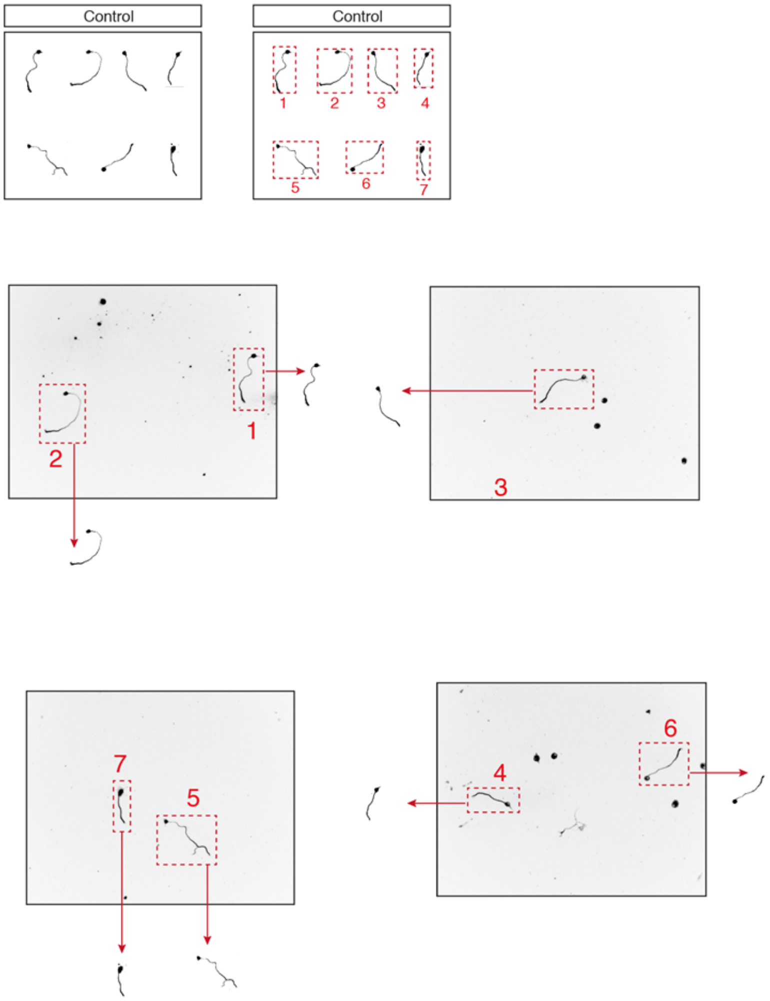


The original microscopic field raw images for constructing the collage image of Fig. 4h, control.

**Supplementary Fig. 10.**

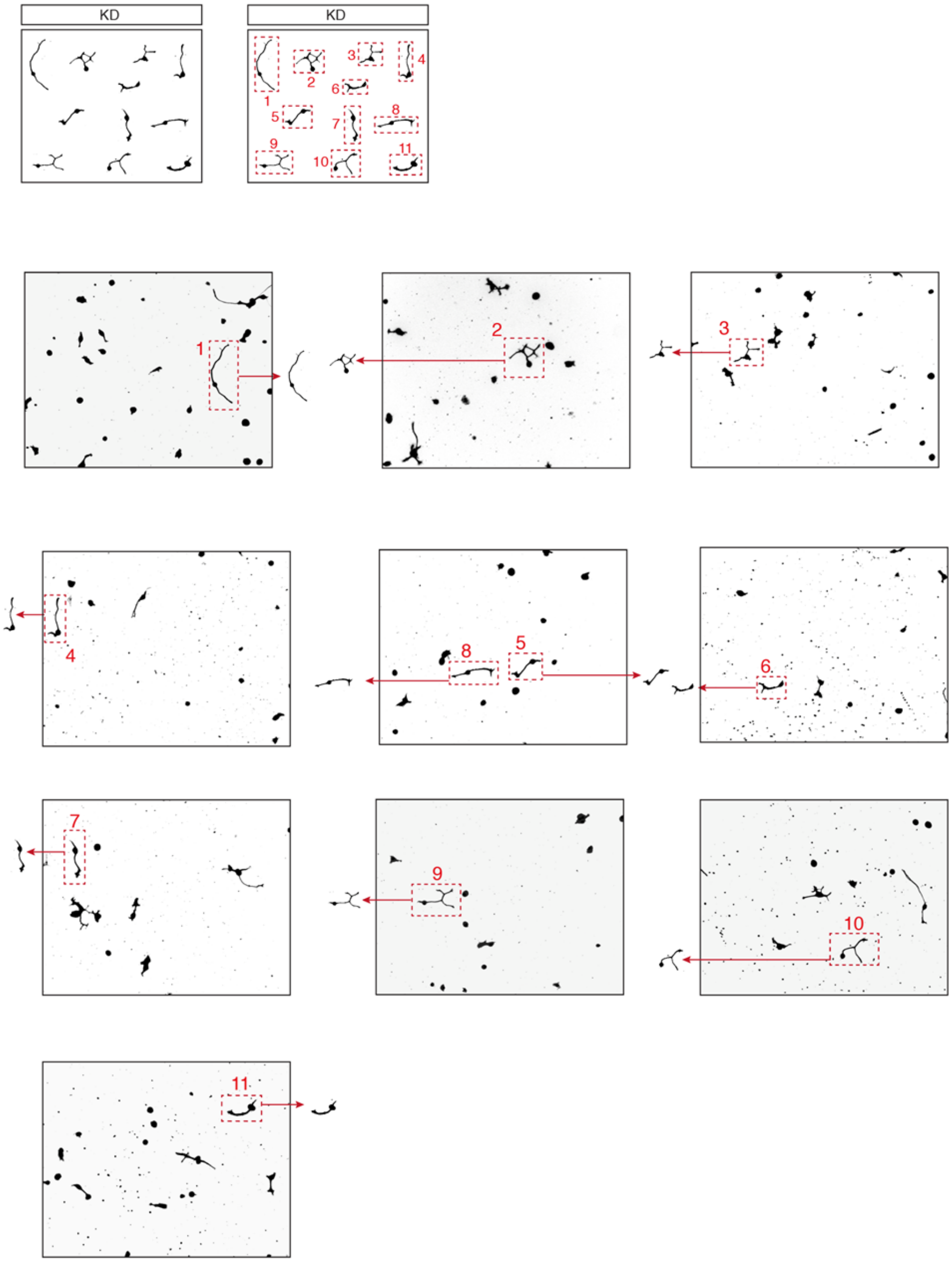


The original microscopic field raw images for constructing the collage image of Fig. 4h, KD.

**Supplementary Fig. 11.**

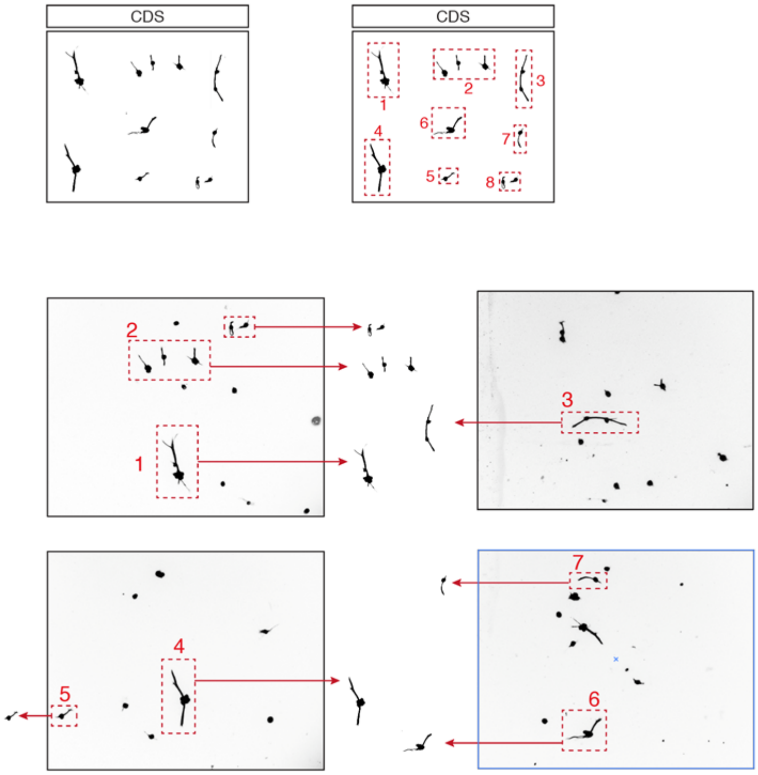


The original microscopic field raw images for constructing the collage image of Fig. 4h, CDS.

**Supplementary Fig. 12.**

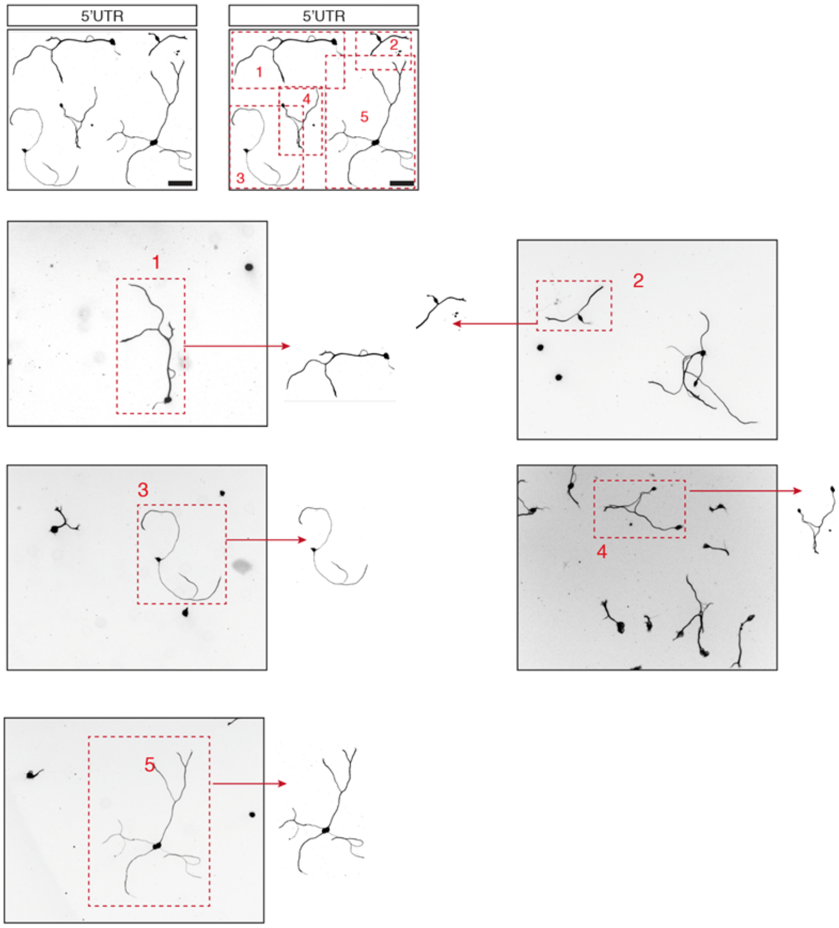


The original microscopic field raw images for constructing the collage image of Fig. 4h, 5'UTR.

**Supplementary Fig. 13.**

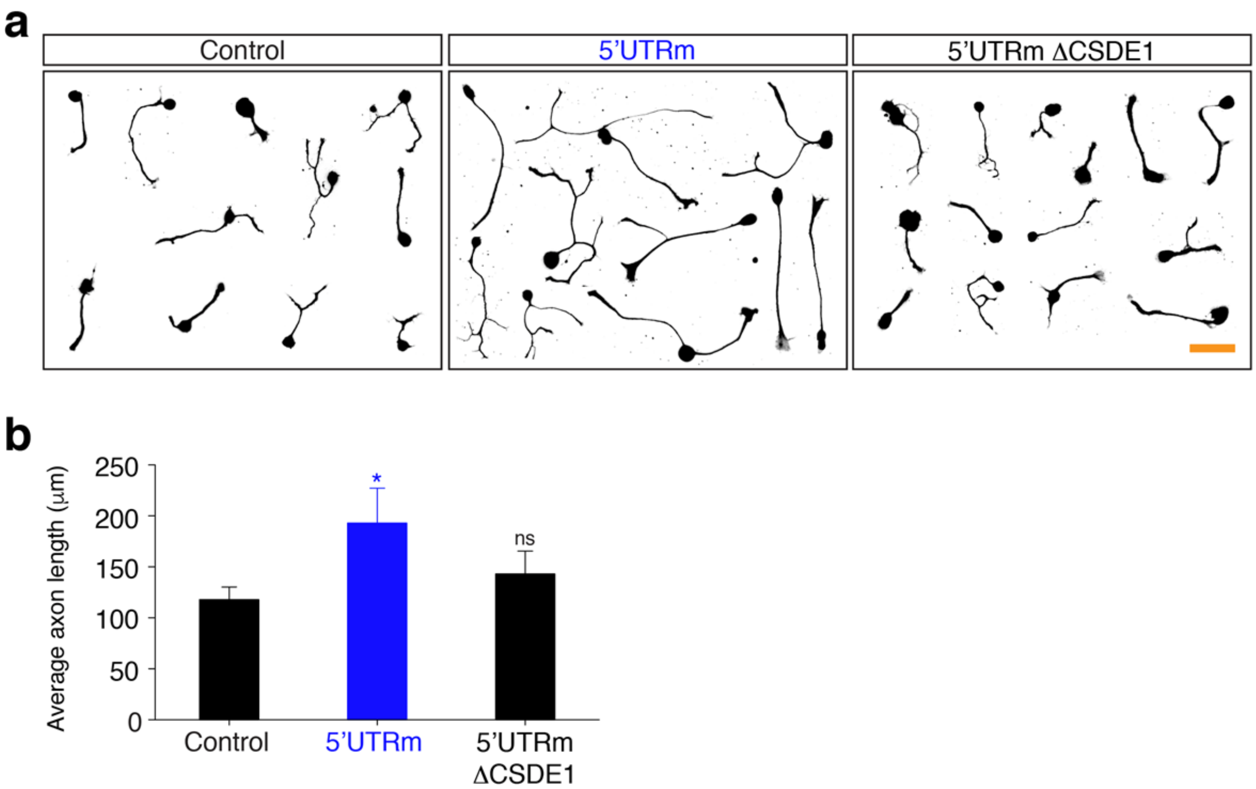


In vitro replating assay of embryonic DRG neurons. (a) Representative images of control, 5’UTRm-overexpressing, and 5’UTRmΔCSDE1-overexpressing embryonic DRG neurons. Scale bar, 100 μm. (b) Statistical analysis of regenerating neurite length of (a) (n=254, 198, 167 for control, shCSDE1 and shKHDRBS1; ***p<0.001, ns, not significant by ANOVA followed by Tukey tests; mean±SEM).

**Supplementary Fig. 14.**

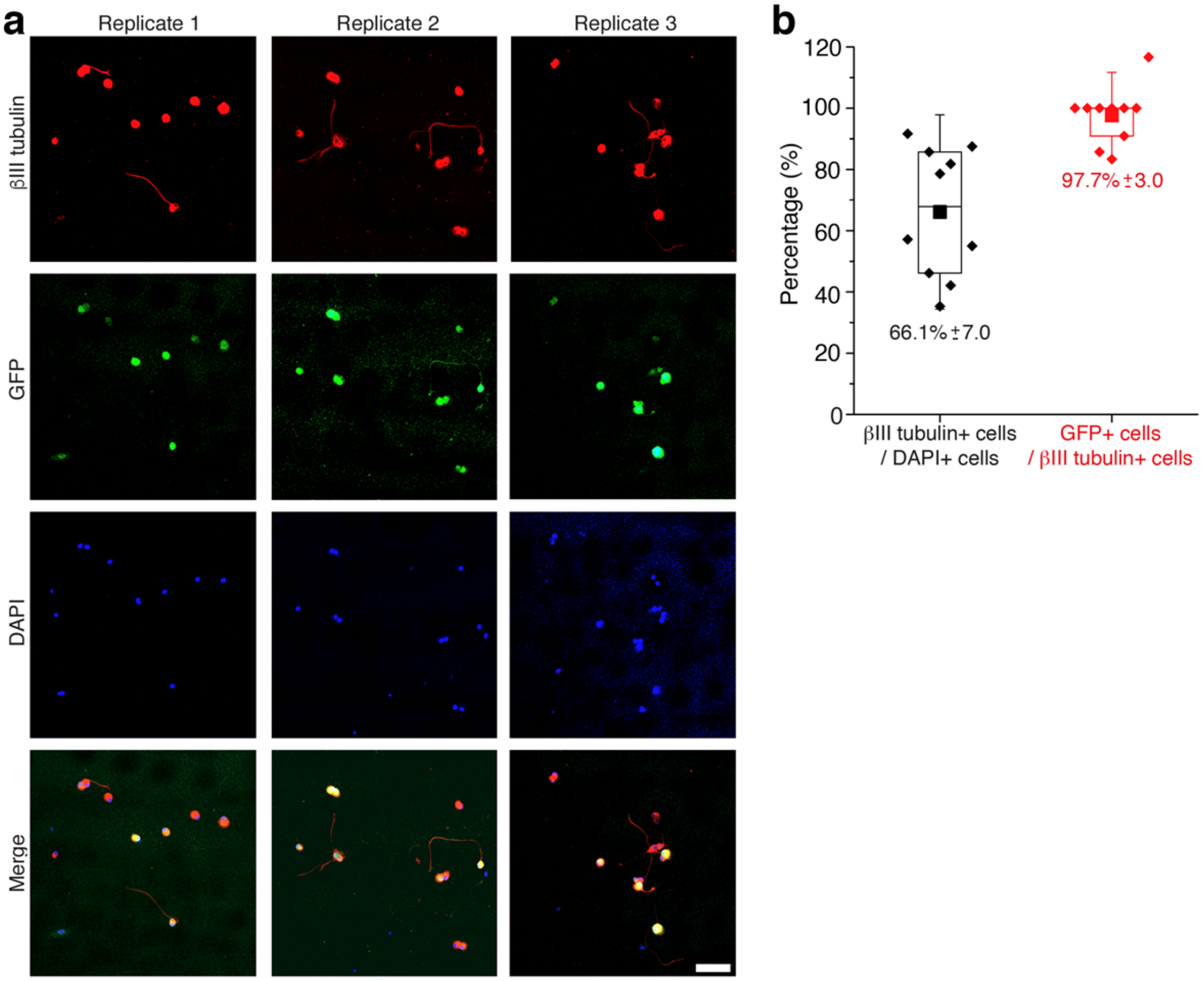


Lentivirus transduction efficiency from in vitro replating assay. Embryonic DRG neurons were prepared by spot-culture method and GFP-expressing control pLKO lentivirus was infected at DIV2. The neurons were replated at DIV5 and fixed and immunostained with βIII tubulin antibody. (a) Representative images of replated DRG neurons from three biological replicates. GFP fluorescence expressed under CMV-promoter of the 2nd ORF of lentivirus vector was visualized. Scale bar, 100 µm. (b) Statistical analysis of the cell numbers from (a). Box indicated 25% to 75% of the distribution. Closed box indicated mean.
